# A Novel Mechanism for NF-κB-Activation via IκB-Aggregation: Implications for Hepatic Mallory-denk-body Induced Inflammation

**DOI:** 10.1101/585497

**Authors:** Yi Liu, Michael J. Trnka, Shenheng Guan, Doyoung Kwon, Do-Hyung Kim, J-J. Chen, Peter A. Greer, A. L. Burlingame, Maria Almira Correia

**Affiliations:** Departments of Cellular & Molecular Pharmacology, University of California San Francisco, San Francisco CA 94158-2517; Pharmaceutical Chemistry, University of California San Francisco, San Francisco CA 94158-2517; Departments of Biochemistry, Molecular Biology, and Biophysics, University of Minnesota, Minneapolis, MN55455; Institute for Medical Engineering and Science, MIT, Cambridge, MA, 02139; Department of Pathology and Molecular Medicine, Queen’s University; Kingston, ON; K7L 3N6; Bioengineering and Therapeutic Sciences, University of California San Francisco, San Francisco CA 94158-2517; The Liver Center, University of California San Francisco, San Francisco CA 94158-2517

**Keywords:** Mallory-Denk-bodies, IκBα, IκBβ, NF-κB, p62, erythropoietic protoporphyria, X-linked protoporphyria, liver inflammation, NMPP, PPIX, ZnPPIX, proteomics

## Abstract

**Background & Aims:** Mallory-Denk-bodies (MDBs) are hepatic protein aggregates associated with inflammation both clinically and in MDB-inducing models. Similar protein aggregation in neurodegenerative diseases also triggers inflammation and NF-κB activation. However, the precise mechanism that links protein aggregation to NFκB-activation and inflammatory response remains unclear.

**Methods:** Herein, we find that treating primary hepatocytes with MDB-inducing agents (N-methylprotoporphyrin, protoporphyrin IX (PPIX), or ZnPPIX) elicited an IκBα-loss with consequent NF-κB activation. We characterized the underlying mechanism in detail using hepatocytes from various knockout mice and MEF cell lines and multiple approaches including immunoblotting, EMSA, RT-PCR, confocal immunofluorescence microscopy, affinity immunoprecipitation, and protein solubility assays. Additionally, we performed rigorous proteomic analyses to identify the proteins aggregating upon PPIX treatment and/or co-aggregating with IκBα.

**Results:** Four known mechanisms of IκBα-loss were probed and excluded. Immunofluorescence analyses of ZnPPIX-treated cells coupled with 8 M urea/CHAPS-extraction revealed that this IκBα-loss was due to its sequestration along with IκBβ into insoluble aggregates. Through proteomic analyses we identified 47 aggregation-prone proteins that co-aggregate with IκBα through direct interaction or proximity. Of these ZnPPIX-aggregation targets, the nucleoporins Nup153 and Nup358/RanBP2 were identified through RNA-interference, as likely mediators of IκBα-nuclear import.

**Conclusion:** We discovered a novel mechanism of inflammatory NF-κB activation through IκB-sequestration into insoluble aggregates along with interacting aggregation-prone proteins. This mechanism may account for the protein aggregate-induced inflammation observed in MDB-associated liver diseases, thereby identifying novel targets for therapeutic intervention. Because of inherent commonalities this MDB cell model is a *bona fide* protoporphyric model, making these findings equally relevant to the liver inflammation associated with clinical protoporphyria.

**Lay Summary:** Mallory-Denk-bodies (MDBs) are hepatic protein aggregates commonly featured in many liver diseases. MDB-presence is associated with the induction of inflammatory responses both clinically and in all MDB-inducing models. Similar protein aggregation in neurodegenerative diseases is also known to trigger inflammation and NFκB pathway activation via an as yet to be characterized non-canonical mechanism. Herein using a MDB-inducing cell model, we uncovered a novel mechanism for NFκB activation via cytosolic IκB-sequestration into insoluble aggregates. Furthermore, using a proteomic approach, we identified 47 aggregation-prone proteins that interact and co-aggregate with IκBα. This novel mechanism may account for the protein aggregate-induced inflammation observed in liver diseases, thereby identifying novel targets for therapeutic intervention.

## INTRODUCTION

Protein aggregates and inclusion bodies are linked to various neurodegenerative, muscular and hepatic diseases [Alzheimer’s, Parkinson’s, Desmin-related myopathies, and Mallory-Denk-bodies (MDBs) [1]]. Although different aggregates vary with each tissue source and in predominant protein composition, they contain common components such as p62/Sequestosome-1, ubiquitin, chaperones and proteasome constituents, which are often misfolded, highly insoluble, and cross-linked [1, 2].

Hepatic MDBs consisting largely of phosphorylated cytokeratins and p62 are commonly found in patients with alcoholic steatohepatitis (ASH) and non-alcoholic steatohepatitis (NASH), primary biliary cirrhosis, non-alcoholic cirrhosis, hepatocellular carcinoma, morbid obesity, and copper-related disorders [3]. MDBs can be reproduced in mice by long term feeding of griseofulvin (GF) or 3,5-dicarbethoxy-1,4-dihydrocollidine (DDC) [4, 5]. Both compounds inactivate certain hepatic cytochromes P450, converting their prosthetic heme into N-methylprotoporphyrin(s) (NMPP) that inhibit hepatic ferrochelatase (Fech) resulting in hepatic heme depletion and protoporphyrin IX (PPIX) accumulation [6, 7] (Fig. 1A). Fech^m1Pas^ mutant (*fech*/*fech*) mice with <5% of normal enzyme activity also develop spontaneous MDBs at 20-weeks of age, validating their use as an experimental model for MDB induction [8].

**FIGURE 1.**
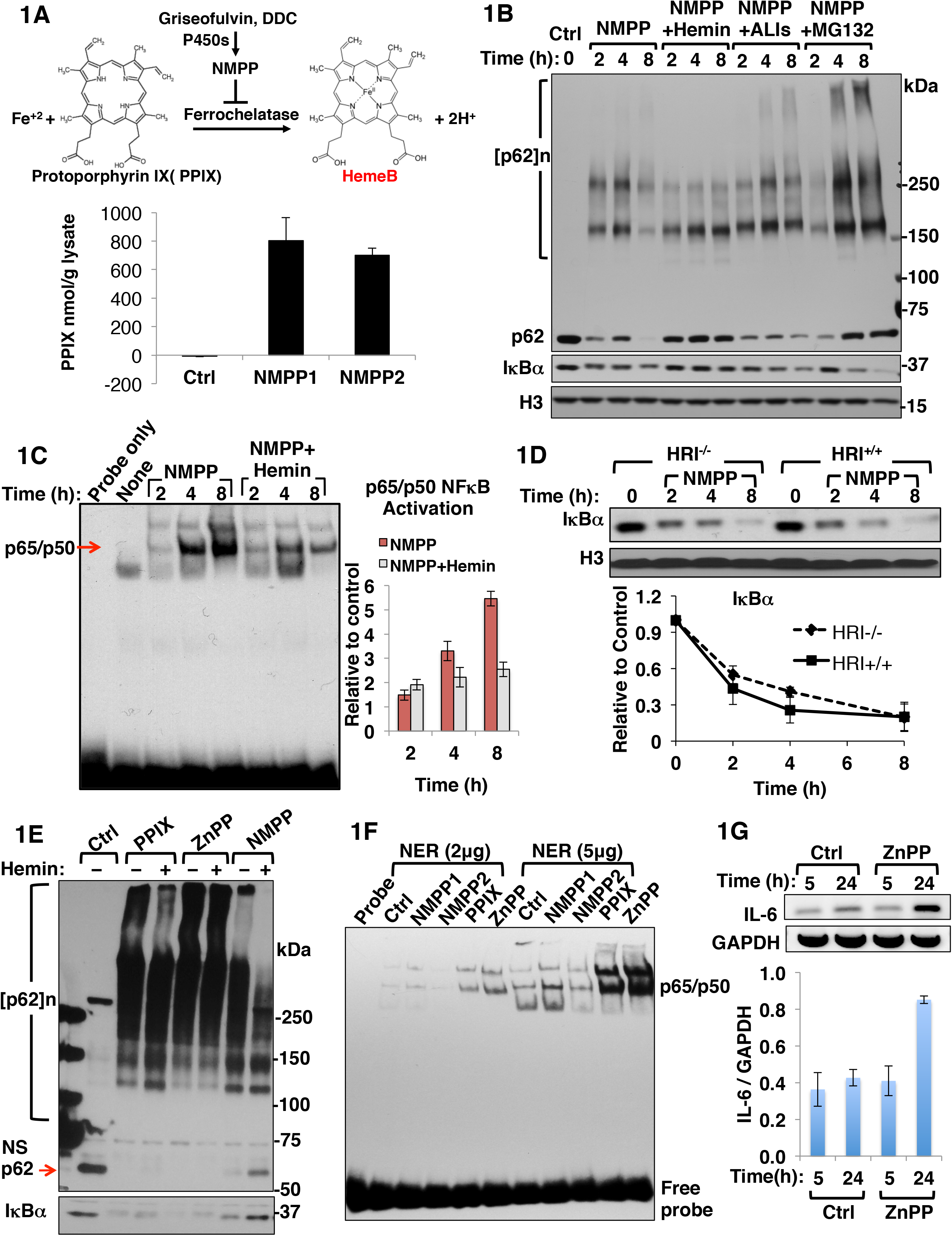
NMPP treatment results in concurrent PPIX-accumulation, NF-κB activation and IκBα-loss. **A**. A scheme for NMPP-mediated inhibition of ferrochelatase with consequent accumulation of the heme precursor PPIX. The PPIX content of lysates from mouse hepatocytes treated with two different commercial lots of NMPP for 24 h quantified flurometrically (Mean ± SD, n = 3). **B**. IB analyses of p62 and IκBα with histone H3 as the loading control in lysates from mouse hepatocytes treated as indicated. NMPP (30 μM), hemin (10 μM added fresh every 4 h), and ALIs, autophagy/lysosomal inhibitors 3MA (5 mM) + NH_4_Cl (30 mM), and MG132 (20 μM) for 1 h. **C**. Mouse hepatocytes were treated as indicated. Representative EMSA of nuclear extracts (NER, 5 μg) with NF-κB-specific oligonucleotides is shown. Bar chart represents mean ± SD of NF-κB activation relative to controls, n = 3. **D**. Quantification of IκBα levels upon IB analyses of lysates from wild type (WT) and HRI^-/-^ mouse hepatocytes treated with 30 μM NMPP for the indicated times (Mean ± SD, n = 3). **E**. IB analyses of p62 and IκBα in lysates from mouse hepatocytes treated with 10 μM PPIX, 10 μM ZnPP or 30 μM NMPP with or without 10 μM hemin for 8 h. NS, non-specific band. **F**. EMSA of nuclear extracts (NER) from mouse hepatocytes treated as indicated for 24 h. **G**. IL-6 RT-PCR analyses of mRNA from mouse hepatocytes treated with 10 μM ZnPP for the indicated times (Mean ± SD, n = 3).

ASH and NASH patient liver histology reveals that MDBs are associated with inflammatory responses, with MDB-containing hepatocytes often surrounded by neutrophils [1, 9]. All MDB-inducing mouse models also accumulate hepatic PPIX and exhibit liver inflammation and injury at an early stage when MDBs are not yet visible [10, 11]. Cumulative biochemical evidence indicates that in these mouse models as well as in DDC-, NMPP- or PPIX-treated cell models, certain proteins such as p62, cytokeratins CK8/18 and lamin begin to aggregate at a much earlier stage before MDBs are detected [12, 13]. Similarly, many neurodegenerative diseases are also associated with inflammation at an early stage, and amyloid protein aggregation has been shown to initiate an inflammatory response [14]. Moreover, induction of various protein aggregates, as in myofibrillar myopathy and Amyotrophic Lateral Sclerosis (ALS), triggers the activation of NF-κB (nuclear factor kappa-light-chain-enhancer of activated B cells) via a noncanonical pathway independent of IκBα-phosphorylation/degradation [15]. However, the precise molecular link between protein aggregation and NF-κB-activation and inflammatory response remains unclear.

Of the existing NF-κB/Rel transcription factors [16], p65/p50 is not only the most ubiquitous and biologically active NF-κB-heterodimer, but also the major hepatic species [17]. p65/p50 is normally sequestered in the cytoplasm by NF-kappa-B inhibitors (IκB). Of these, in human hepatocytes, HepG2 cells and cultured mouse hepatocytes, IκBα and IκBβ are the major forms, each regulated by different signals [18-21]. IκB-binding masks NF-κB nuclear localization signal (NLS) and DNA-binding domain [16]. Signal-induced IκB-unleashing of cytoplasmic NF-κB most commonly via phosphorylation and subsequent ubiquitin-dependent degradation (UPD) of both IκBα and IκBβ [22-24], results in the nuclear translocation of DNA-binding competent NF-κB, with consequent transcriptional activation of its target genes including that of IκBα [25], but not of IκBβ [26].. IκBα is then rapidly *de novo* oversynthesized (overshoot), and following nuclear import, binds and displaces the DNA-bound NF-κB, masking its NLS and accelerating its nuclear export, required for down regulation and eventual termination of NF-κB-mediated transcriptional activation [27]. Thus, while both hepatic IκBs are largely involved in NF-κB-cytoplasmic retention, only IκBα is involved in nuclear NF-κB-transcriptional suppression [19]

Our findings herein reveal a loss of IκBα and IκBβ with subsequent NF-κB activation upon treatment of primary hepatocytes with NMPP (an established MDB inducer), or ZnPPIX (ZnPP). We probed the role of four plausible mechanisms for NF-κB-activation via reduction of intracellular IκBα-levels in this NMPP-elicited IκBα-loss and excluded them all (Supplementary Results). Instead, we found that this apparent IκBα-loss upon NMPP- or ZnPP-treatment is actually due to its phase sequestration into insoluble cellular aggregates along with IκBβ, as well as the nucleoporins Nup153 and Nup358 that are involved in IκBα nuclear import and subsequent termination of NF-κB activation.

To our knowledge, this is the first evidence that NF-κB is also activated through IκB-sequestration into insoluble aggregates. We believe such a novel mechanism accounts for the persistent hepatic NF-κB activation that may directly contribute to the severe inflammatory responses and liver injury observed not only in various MDB-featuring liver diseases and experimental MDB-models but also in acute erythropoietic protoporphyria (EPP) and X-linked protoporphyria (XLPP) [28-30].

## RESULTS

### NMPP-elicited PPIX accumulation with concurrent IκBα-loss and NF-κB activation in cultured mouse hepatocytes

NMPP-treatment of cultured mouse hepatocytes as expected from its ferrochelatase inhibition, resulted in cellular PPIX accumulation (Fig. 1A). In parallel, a gradual IκBα loss was also observed in hepatic lysates upon NMPP-treatment (Fig. 1B). This IκBα-loss was reversed upon inclusion of hemin, but not that of the proteasomal inhibitor MG-132, or the dual autophagic-lysosomal degradation (ALD) inhibitors 3-methyladenine (3MA) + NH_4_Cl. These findings thus excluded both UPD and ALD involvement in this IκBα-loss. Further supportive evidence was provided by studies in HepG2 cells transfected with a S_32_A/S_36_A-IκBα mutant that is resistant to phosphorylation and UPD (Fig. S1A) as NMPP caused a similar time-dependent loss of both WT IκBα and its mutant. Parallel EMSA of nuclear extracts from these NMPP-treated hepatocytes revealed a time-dependent hepatic NF-κB activation, indicating that his hepatic IκBα loss was indeed physiologically relevant (Figs. 1C & S1B). This NF-κB activation was also attenuated by hemin (Fig. 1C), consistent with the hemin-mediated inhibition of IκBα loss (Fig. 1B). These findings suggested that this culture model was a valid model to interrogate the association of PPIX accumulation with NF-κB-activation and inflammation.

Four additional mechanisms -- two established and two plausible -- for NF-κB-activation via reduction of intracellular IκBα-levels [i.e. (i) translational suppression by heme-deficiency-triggered heme-regulated inhibitor (HRI) eIF2α kinase-activation ([31]; Figs. 1D & S2A); (ii) enhanced autophagy ([32]; Fig. S2B); (iii) enhanced calpain-mediated proteolysis ([33]; Fig.S2C); and (iv) PPIX-photoactivation and consequent ROS-mediated oxidative stress [28] in NMPP-elicited IκBα-loss (Supplementary Results; Fig. S3), were all probed and excluded.

### PPIX and ZnPP are even more potent inducers of IκBα-loss, NF-κB-activation and p62 oligomerization/aggregation than NMPP

Because NMPP-elicited ferrochelatase inhibition results in PPIX-accumulation, we examined whether this accumulation (and not heme deficiency) was mainly responsible for the observed IκBα loss and NF-κB activation. We observed a greater IκBα loss after either PPIX or ZnPP relative to that seen with NMPP alone (Fig. 1E). While the NMPP-elicited IκBα loss was to a great extent prevented by hemin inclusion, the IκBα loss after PPIX or ZnPP was not (Fig. 1E; see Supplementary Discussion). EMSA analyses revealed a NF-κB activation that was proportional to the observed IκBα loss, ranking in order as follows: ZnPP>PPIX>NMPP (Fig. 1F). Consistent with this NF-κB activation, RT-PCR analyses of mRNA from ZnPP-treated hepatocytes indicated marked up-regulation of the inflammatory cytokine IL-6 at 24 h (Fig.1G).

Because p62 is a major component of MDB and various other pathological aggregates [1], we monitored hepatic p62 response upon NMPP, PPIX and ZnPP ± hemin treatments (Figs. 1B, 1E). In parallel with IκBα loss, p62 formed aggregates and possibly cross-linked species, with a concurrent decline in the monomeric species (∼62 kD; Ctrl), evident at later (8 h) rather than earlier (2-4 h) time points. This apparent time-lag is most likely due to an initial counteractive compensatory p62 transcriptional induction in response to PPIX-elicited oxidative stress and Nrf2 activation [8]. Furthermore, we found that the p62-monomer levels correlated reasonably well (r^2^ = 0.72) with corresponding IκBα levels in these treated and untreated cells, revealing an intimate association between IκBα loss and p62-aggregation, possibly due to their direct protein interactions (Fig. S1D).

### ZnPP-elicited IκBα-loss is independent of p62 or other autophagic receptors/adapters

The capacity of p62 to stabilize IκBα upon co-expression in HEK293T cells (Fig. 2A) and the close temporal relationship between IκBα-loss and p62 aggregation (Fig. 1B) suggested these proteins were intimately associated. We thus examined their interdependence in ZnPP-treated p62 WT and p62 null mouse (p62KO) hepatocytes or MEF cells (Figs. 2B & S4A), and found that this IκBα loss was independent of p62 aggregation. The additional possibility that in p62-lacking cells, IκBα loss was due to its degradation via UPD, ALD or calpain pathways was excluded with specific inhibitors of these pathways (Fig. S4B). We also examined plausible compensation by NBR1 (Neighbour of Braca 1 gene) [34] (Fig S4C, S4D), and three other autophagic receptors/adapters [35], and conclusively excluded p62 and its redundant functional mimics in this ZnPP-elicited IκBα-loss (Fig. 2D).

**FIGURE 2.**
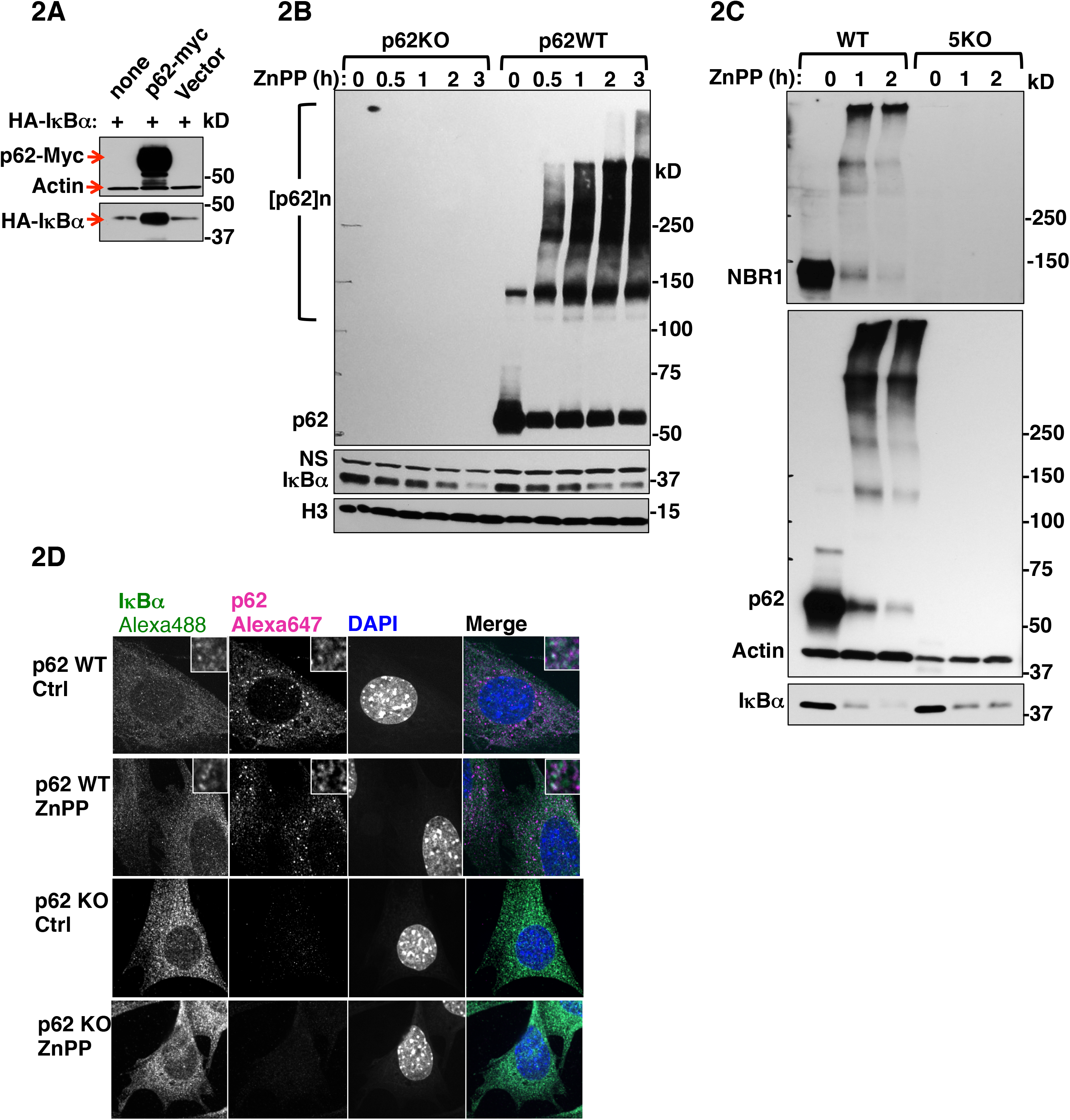
ZnPP-elicited IκBα-loss is p62-independent. **A**. HEK293T cells were transfected with pCMV4-3HA-IκBα, or co-transfected with either pcDNA6-p62-Myc or pcDNA6-Myc empty vector for 48 h. Cell lysates were used for IB analyses with actin as the loading control. **B**. Primary hepatocytes from wild type (p62WT) and p62 knockout (p62KO) mice were treated with 10 μM ZnPP for the indicated times. Cell lysates were used for IB analyses. **C**. Wild-type Hela cells (WT) and CRISPR-engineered Hela cells with five autophagic adapters genetically deleted (5KO) were treated with 10 μM ZnPP for the indicated times. Lysates were used for IB analyses. **D**. CMIF analyses. p62WT and p62KO MEF cells were treated with 10 μM ZnPP or vehicle control for 2 h, and then fixed and stained with anti-IκBα (green), anti-p62 (magenta) and DAPI (Blue). Insets depict enlarged regions of IκBα and p62 colocalization.

### Cytoplasmic IκBα-loss is due to its physical sequestration into insoluble cellular aggregates

Confocal microscopic immunofluorescence (CMIF) analyses of ZnPP-treated WT and p62^-/-^-MEF cells provided an informative clue (Fig. 2E). In WT-cells, both proteins were largely colocalized to the cytoplasm, and ZnPP did not affect this colocalization (Fig. 2E, insets). Surprisingly, following ZnPP-treatment, in spite of immunochemical IκBα-loss (Fig. 2D), IκBα-associated immunofluorescence signal still persisted comparably in both WT and p62^-/-^-MEF cells. This finding provided the first clue that upon ZnPP-treatment, IκBα was not irretrievably lost, but just undetectable in detergent-solubilized cell lysates routinely employed for immunoblotting (IB) analyses. It was thus likely that such an IκBα physical inaccessibility was due to intracellular ZnPP-triggered protein aggregation. To examine this possibility, HEK293T cells co-transfected with both HA-IκBα and p62-Myc plasmids, were treated with ZnPP and then sequentially extracted using detergents of increasing strengths (Supplementary Methods). IB analyses showed that upon ZnPP-treatment, in parallel with p62 aggregation, HA-IκBα became undetectable in soluble Triton and RIPA fractions (Fig. 3A), however, the majority of HA-IκBα as well as monomeric p62 and p62-aggregates could be recovered from the RIPA-insoluble pellet by heat extraction with 8M urea/CHAPS buffer (urea buffer; Supplementary Methods). Furthermore, similar results were found upon IB analyses of these urea extracts irrespective of whether antibodies to the HA-tag, N- or C-terminus of IκBα were employed (Fig. 3B). IB analyses of high salt buffer (HSB)-extracts from ZnPP-treated cell lysates also revealed detectable monomeric and aggregated IκBα and p62 species, albeit to a much lesser extent than corresponding urea extracts (Fig. 3C). Together, these findings indicated that upon ZnPP-treatment, p62 and IκBα along with other cellular proteins co-aggregate, and these aggregates are sequestered from the cytoplasm into an insoluble cellular fraction, essentially unavailable to function physiologically.

**FIGURE 3.**
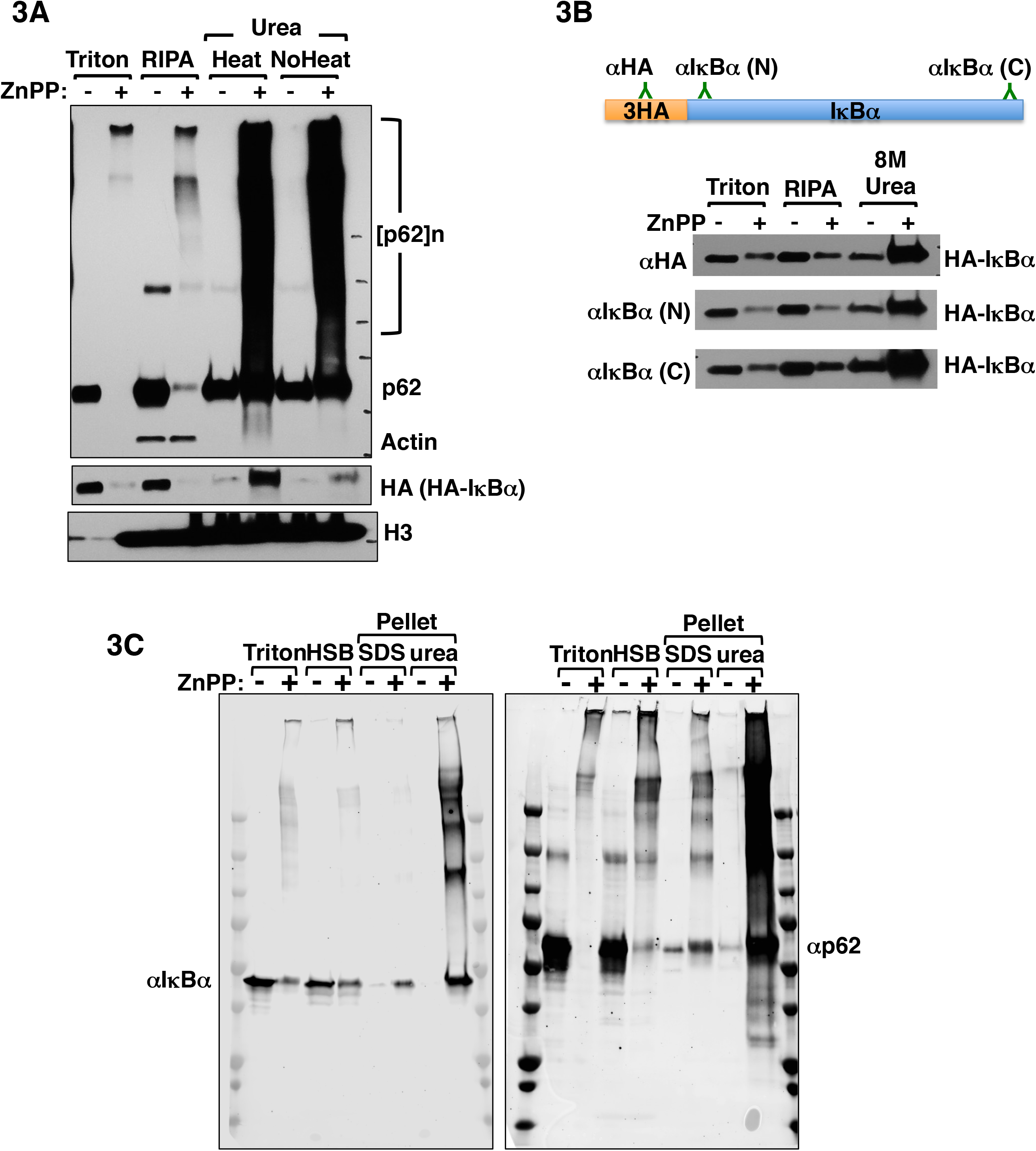
ZnPP-elicited IκBα-loss is due to its sequestration into insoluble cellular aggregates. **A**. HEK293T cells were transfected with pCMV4-3HA-IκBα for 40 h, and then treated with 10 μM ZnPP for 4 h. Cells were then subjected to sequential extraction. 10 µg of extracts were used for IB analyses. **B**. The same extracts from (**A**) were immunoblotted using three different antibodies each targeting a different region of the same HA-IκBα protein: Anti-HA-tag, rabbit monoclonal antibody targeting IκBα N-terminus (N), and rabbit monoclonal antibody targeting IκBα C-terminus (C). **C**. HEK293T cells were transfected and treated as in **A**. and then subjected to sequential extraction with the following buffers: Cell Signaling lysis buffer (Triton), Cell Signaling lysis buffer supplemented with 1.5 M KCl (HSB). The pellet obtained after HSB-extract was either solubilized with heating in a Laemmli buffer with 4% SDS (SDS) or in urea buffer. 10 µg of extracts were used for IB analysis.

### Proteomic identification of plausible IκBα-interactors and ZnPP-aggregated proteins

PPIX-treatment is causally associated with the highly selective aggregation of many cellular proteins [36]. Thus, IκBα could either be the prime target of ZnPP-elicited physical sequestration or a silent partner dragged along for the ride by one or more of its cellular interactors. To identify any plausible mediators of ZnPP-elicited IκBα sequestration into insoluble cellular aggregates, we employed a proteomic approach to comprehensively identify the proteins that normally interact with IκBα, but which form protein co-aggregates upon ZnPP-treatment. These aggregation-prone proteins upon IκBα-interaction could drive its sequestration and co-aggregation.

To obtain a high-confidence IκBα-interactome, we performed rigorous immunoaffinity purification coupled with mass spectrometry (IAP-MS) analyses, with extensive biological repetitions and strict filtering (Fig. 4A, 4B, Supplementary Methods). This approach identified 370 high-confidence IκBα-interactors, including well-established IκBα-interactors [i.e. NF-κB subunits (RelA, NFKB1, NFKB2), and IKK-complex proteins (CHUK, IKBKAP)] (Fig. 4C), thus validating our IAP-MS approach. However, most of the proteins identified were novel IκBα-interactors.

**FIGURE 4.**
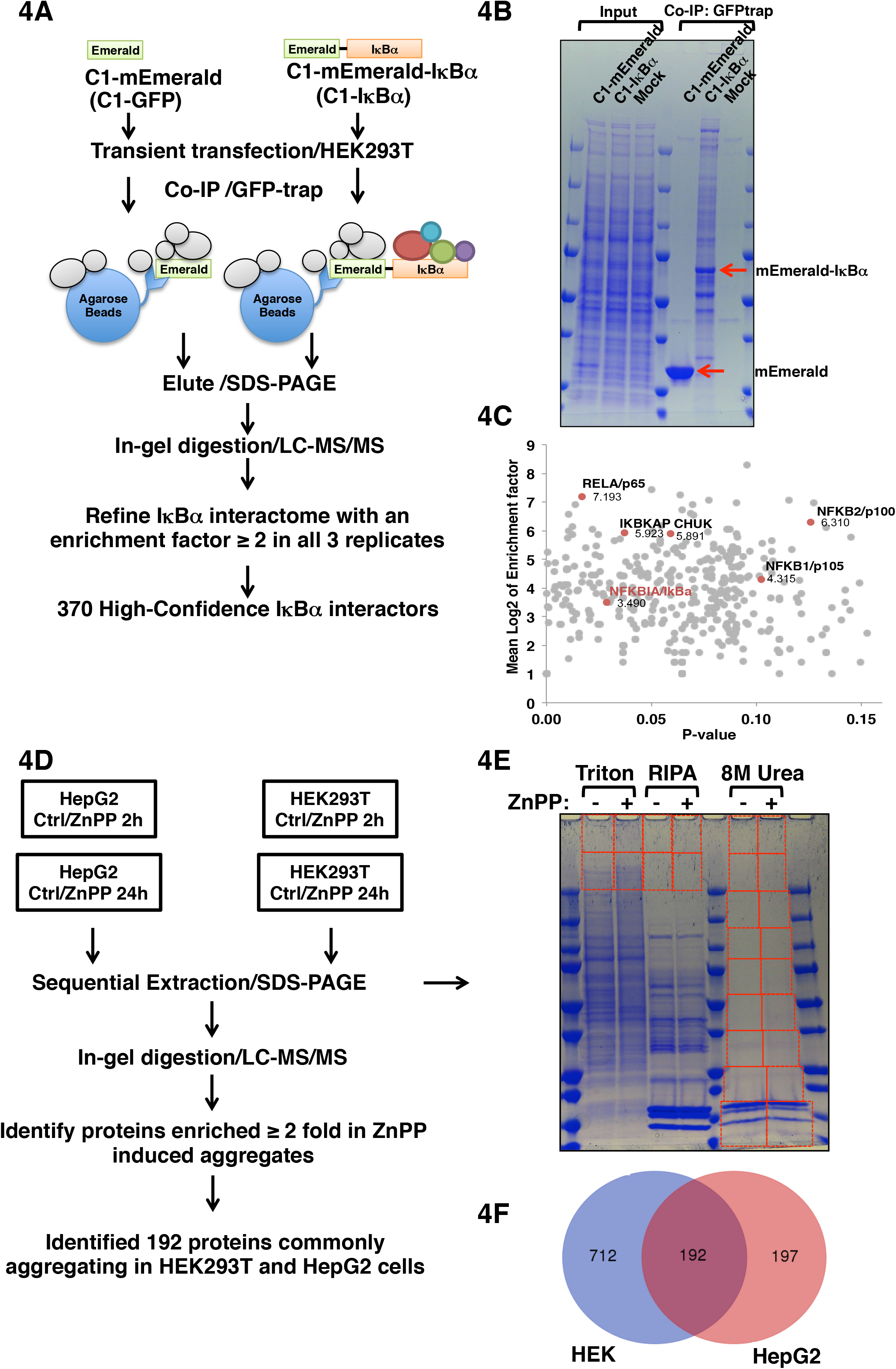
IAP-MS of IκBα-interactome and ZnPP-induced aggregate proteome. **A**. Flow-chart of the strategy used to identify a high-confidence IκBα-interactome. Cells expressing GFP (mEmerald) alone were used as the background control. **B**. Representative Coomassie Blue staining lysates before and after IAP. **C**. 370 high-confidence IκBα-interactors were plotted with the mean log of enrichment factor against GFP-control on the Y-axis and the *p*-value on the X-axis. Previously known IκBα**-**interactors are labeled in red. **D**. Flow-chart of the strategy employed to identify ZnPP-induced aggregate proteome of both HepG2 and HEK293T cells. **E**. Coomassie Blue staining of proteins from sequential Triton, RIPA and urea fractions. HMM fractions of Triton and RIPA extracts and the entire urea fraction (demarcated by the red boxes) were excised from the gel for in-gel digestion and MS analyses (LC-MS/MS). **F**. Venn diagram of proteins enriched at least 2-fold in ZnPP-induced aggregates relative to corresponding controls from HEK293T (HEK) and HepG2 cells indicated 192 proteins as common hits.

Previous studies in *fech/fech* mouse and protoporphyric zebra fish models have identified several (≈ 30) aggregated proteins in the high molecular mass (HMM)-regions upon SDS-PAGE of NP40-soluble and/or SDS-soluble liver fractions [8, 13, 36, 37]. Our discovery that ZnPP-elicited protein aggregates could be resolubilized by heating in urea buffer, prompted comprehensive proteomic analyses of these urea-solubilized protein aggregates. For this purpose, the progressive temporal course of this protein aggregation process was followed in HEK293T and HepG2 cells treated with ZnPP for 2 h and 24 h (Fig. 4D). The entire urea-solubilized fraction as well as the HMM-bands from SDS-PAGE of Triton and RIPA soluble extracts were subjected to LC-MS/MS proteomic analyses (Fig. 4D, E). Overlap analyses revealed that 192 proteins commonly aggregated in both cell lines upon ZnPP-treatment (Fig. 4F).

### Disruption of multiple functional networks upon ZnPP-elicited physical sequestration of IκBα and other cellular proteins

To gain functional insight into these 192 ZnPP-aggregated proteins and their possible contribution to cell toxicity, we undertook a comprehensive protein network analyses using the Search Tool for the Retrieval of Interacting Genes/Proteins (STRING) coupled with biological process and pathway overrepresentation analyses (Fig. 5A). Clearly, ZnPP-elicited physical aggregation clustered these proteins under 4 critical biological processes. Those proteins occupying a network hub/node position within their individual clusters, could disrupt multiple vital cellular interactions, leading to global hepatic functional collapse. Additional overlap of the 370 IκBα-interacting proteome with the common 192 ZnPP-aggregate proteome identified 47 IκBα-interacting proteins in common with the ZnPP-aggregate proteome(*each highlighted in yellow*; Fig. 5A). By contrast, well-established IκBα interactors (RelA, NFKB1, NFKB2, CHUK and IKBKAP) were not detected within the ZnPP-aggregate proteome of either cell type (Fig. 5A). An enrichment ratio of the 47 overlapping proteins under each experimental condition was then employed to percentile rank them in a heat-map clustering format (Fig. 5B). Such analyses identified many nuclear pore complex (NPC) nucleoporins Nup153, Nup155, and Nup93, and Nup358/SUMO E3-ligase RANBP2, as significantly enriched in this combined proteome (Fig. 5B).

**FIGURE 5.**
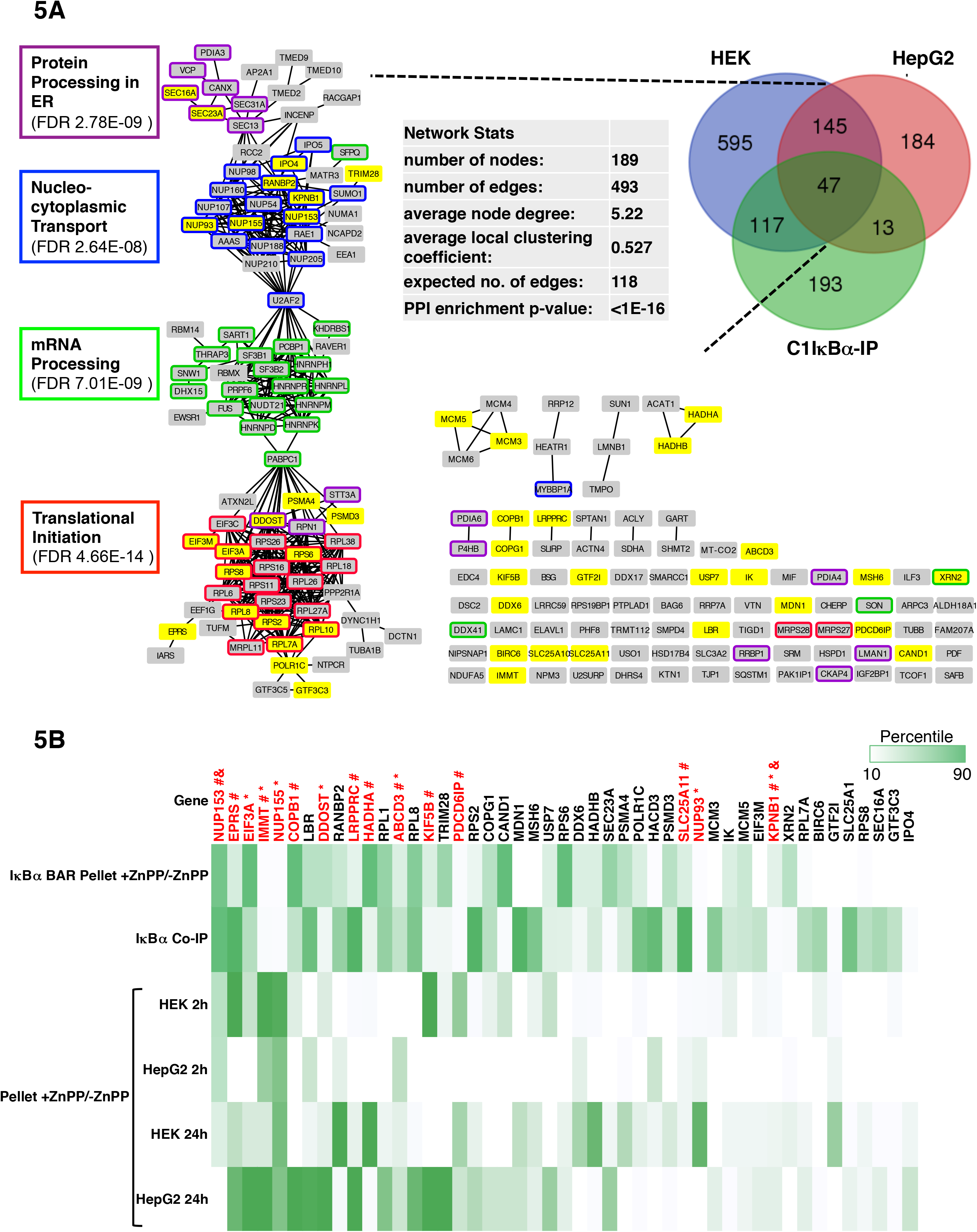
STRING network analyses of proteins common to ZnPP-induced cellular aggregates and IκBα-interactome. **A**. Venn diagram of IκBα**-**interacting proteins identified through LC-MS/MS proteomic analyses of IκBα-immunoprecipitates upon IAP and/or ZnPP-induced cellular aggregates from HEK293T and HepG2 cells (*as schematized in* Fig. 4). The complete interaction network of 192 common ZnPP-induced protein aggregates was obtained via STRING analyses with the highest confidence score (≥ 0.9). Network statistics are summarized in the inset. Functional enrichments are annotated with color-lined boxes with the false discovery rates (FDR) included. Of these, the 47 IκBα**-**interacting proteins are highlighted in yellow. **B**. The corresponding heat map percentile ranked according to the individual fold-enrichment of these 47 IκBα**-**interacting proteins common to ZnPP-induced aggregates in different experiments. C1-IκBα IP represents mean fold-enrichment from C1-mEmerald-IκBα IAP against C1-Emerald (GFP) background control (n = 3). The other sets represent fold-enrichment of proteins in cellular aggregates at 2 h and 24 h post ZnPP-treatment of HepG2 and HEK293T cells relative to corresponding controls. IκBα BAR in +ZnPP/-ZnPP pellet aggregates represents fold-enrichment of IκBα-antibody mediated IP of biotinylated proteins in cellular aggregates at 2 h after ZnPP-treatment of HEK293T cells relative to vehicle controls. Proteins previously found in aggregates from other disease models and/or human pathological samples are indicated in red. * Proteins found in liver aggregates from mouse and zebra fish protoporphyric models [13, 36, 37]. ^#^ Proteins previously found in protein aggregates of pathological samples from human MS and/or Lewy bodies of Parkinson’s disease [53, 55]. ^&^ Proteins previously found in aggregates/ or mislocalized in ALS/FTLD patient samples [54].

To further verify that these 47 proteins were indeed co-aggregating with IκBα, rather than merely aggregating in parallel, we employed biotinylation proximity labeling to identify IκBα-interacting proteins within the protein aggregates. After due consideration of various proximity labeling approaches, we selected HRP-biotinylation by antibody recognition (BAR) [38] for identification of IκBα-interacting proteins in both soluble and insoluble fractions of ZnPP-treated and untreated cells (Fig. S5). The overlapping proteome identified via both co-IP and BAR approaches revealed a very high-confidence IκBα-interactome. More importantly, of the previously identified 47 IκBα-interacting proteins in common with the ZnPP-aggregate proteome, 37 were also detected via BAR to be interacting with IκBα in ZnPP-triggered protein aggregates (Fig 5B), indicating that they were indeed co-aggregating.

### Nucleoporin Nup153 is a principal IκBα-interactant and ZnPP-target

The heat-map clustering analyses of these 47 proteins consistently identified the nuclear pore basket component Nup153 as the top common IκBα-interactant in IAP-MS, BAR, and HEK293 and HepG2 ZnPP-aggregate proteome. We therefore verified Nup153 presence through IB analyses of Triton- and urea-solubilized cellular aggregates upon a 2 h-ZnPP-treatment of HEK293 and HepG2 cells. Indeed, whereas native Nup153 exhibited its intrinsic 153 kDa-mobility in untreated cells, upon ZnPP-treatment it was found as HMM-protein aggregates in Triton- and urea extracts, along with endogenous IκBα (Fig. 6A). Furthermore, co-immunoprecipitation (Co-IP) using HEK293T cells revealed a significant fraction of endogenous Nup153 interacted with IκBα both in cytoplasmic and nuclear extracts of GFP-IκBα-transfected cells, but not in mock- or C1-GFP-transfected cells (Fig. 6B). A similar interaction was also evident among the endogenous counterparts under basal conditions. However, this Nup153-IκBα-interaction was greatly enhanced upon TNFα-treatment at times (1 and 1.5 h) when rapid translocation of *denovo* synthesized IκBα across the nuclear pore would be expected to trigger its post-induction repression of NF-κB-activation (Fig. 6C). Furthermore, ZnPP-treatment of TNFα-pretreated cells led to accelerated IκBα sequestration, whose timing revealed that ZnPP may preferentially target *de novo* synthesized IκBα as it increasingly interacts with cytoplasmic Nup153 during its nuclear import (Fig. 6C-D). Consistent with this, our CMIF of HepG2 cells revealed that under basal conditions, both NF-κB (p65-Rel A subunit) and IκBα are localized in the cytoplasm (Fig. 6E). But upon TNFα-treatment, IκBα UPD results in NF-κB nuclear translocation within 0.5 h. Subsequently, upon transcriptional activation of the NF-κB-responsive IκBα-gene, newly synthesized IκBα enters the nucleus resulting in NF-κB dissociation and nuclear export, all within 1 h of TNFα-treatment. By contrast, upon ZnPP-treatment, in spite of this robust IκBα-restoration at 1 h after TNFα-treatment, it is apparently functionally defective as a nuclear NF-κB-repressor, as NF-κB persisted in the nucleus (Fig. 6E). This stalled IκBα appears prominently both in the cytoplasm as well as clustered around the outer nuclear envelope rim (Fig. 6E). Additional support for this likelihood is provided by our Nup153 and RanBP2 siRNA-knockdown analyses that revealed the marked attenuation of nuclear IκBα-import upon TNFα-activation, without affecting its ZnPP-sequestration (Figs. 7 & S6).

**Figure 6:**
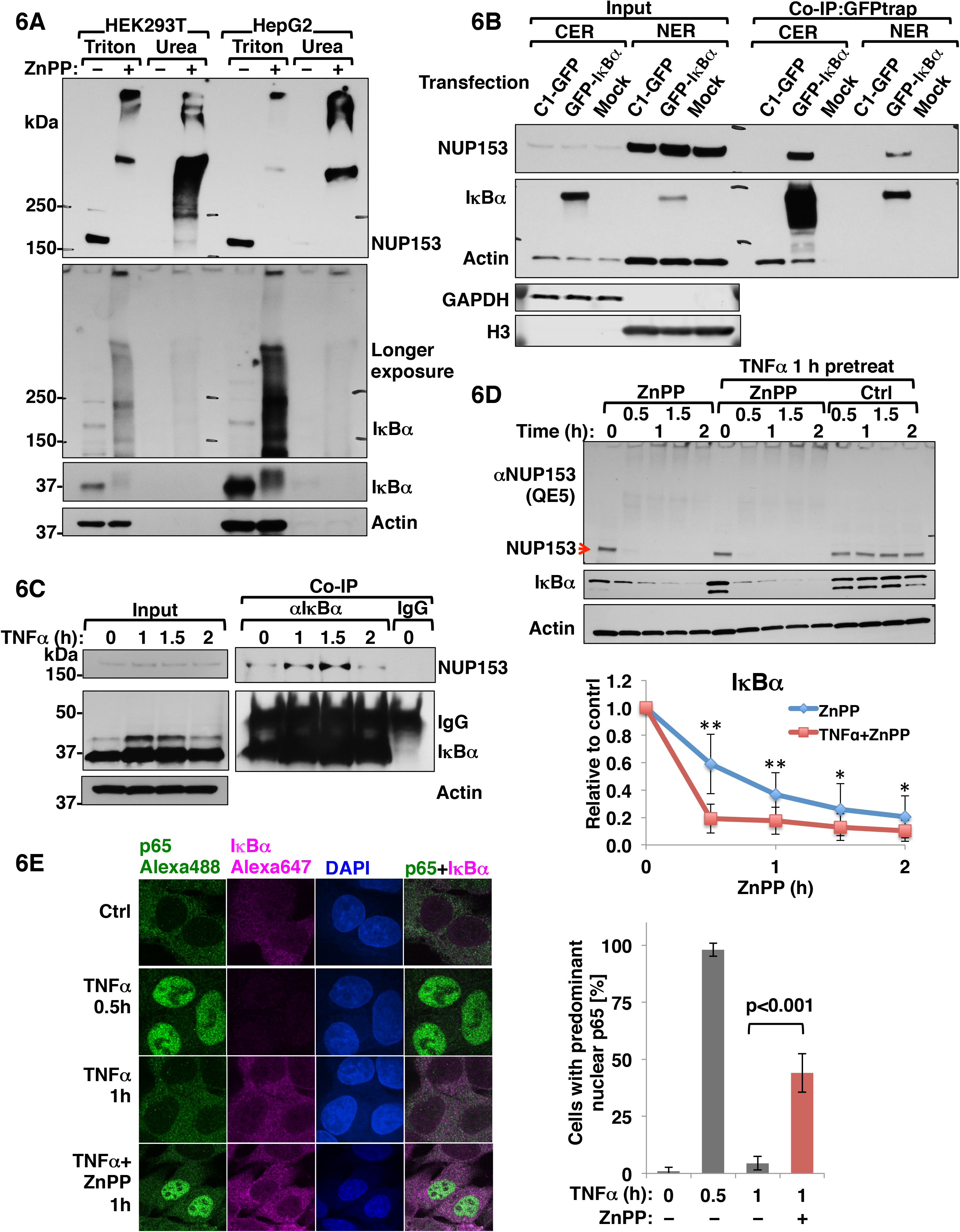
Nup153 is an IκBα-interacting protein that is a major target of ZnPP-elicited aggregation. **A.** HEK293T cells and HepG2 cells were treated with 10 µM ZnPP for 2 h. Cells were sequentially extracted with Triton and urea buffers. 10 µg of extracts were used for Nup153 and IκBα IB analyses, with actin as the loading control. **B.** Co-immunoprecipitation (Co-IP) of overexpressed IκBα with endogenous Nup153. HEK293T cells were transfected with C1-GFP control vector, or C1-GFP-IκBα vector, or a mock control for 40 h. Co-IP of cytosolic fractions (CER) and nuclear fractions (NER) with GFP-trap followed by Nup153 and IκBα IB analyses, with actin as the loading control, and GAPDH and HistoneH3 (H3) as fractionation controls. **C**. Co-IP of endogenous IκBα with endogenous Nup153. HepG2 cells were treated with 20 ng/ml TNFα for the indicated times followed by co-IP of whole cell lysates with IκBα antibody or control IgG and IB of Nup153 and IκBα with actin as the loading control. **D.** Mouse hepatocytes were either untreated or pretreated with 20 ng/ml TNFα for 1 h and then treated with 10 µM ZnPP or left untreated for the indicated times. Cell lysates were used for IB analyses of Nup153 and IκBα with actin as the loading control. IκBα contents relative to 0 h controls were quantified (Mean ± SD, n = 3, ** P<0.01, * P<0.05). **E.** CMIF analyses. HepG2 cells were treated with vehicle control (Ctrl), or 20 ng/ml TNFα for 0.5 h or 1 h. Some TNFα-treated cells were also treated with 10 µM ZnPP 20 min after TNFα-addition and incubated for another 40 min (TNFα + ZnPP 1h). Cells were fixed and stained with anti-p65/anti-rabbit-Alexa-488 IgGs (green) and anti-IκBα/anti-mouse Alexa-647-conjugated IgGs (magenta) and DAPI. Images were obtained under 100X lens. The same slides were then observed under wide-field microscope using 60X lens to quantify the percentage of cells with predominant nuclear p65 accumulation. Cells (>600) were assessed at each experimental condition (Mean ± SD, n = 3).

**Fig. 7.**
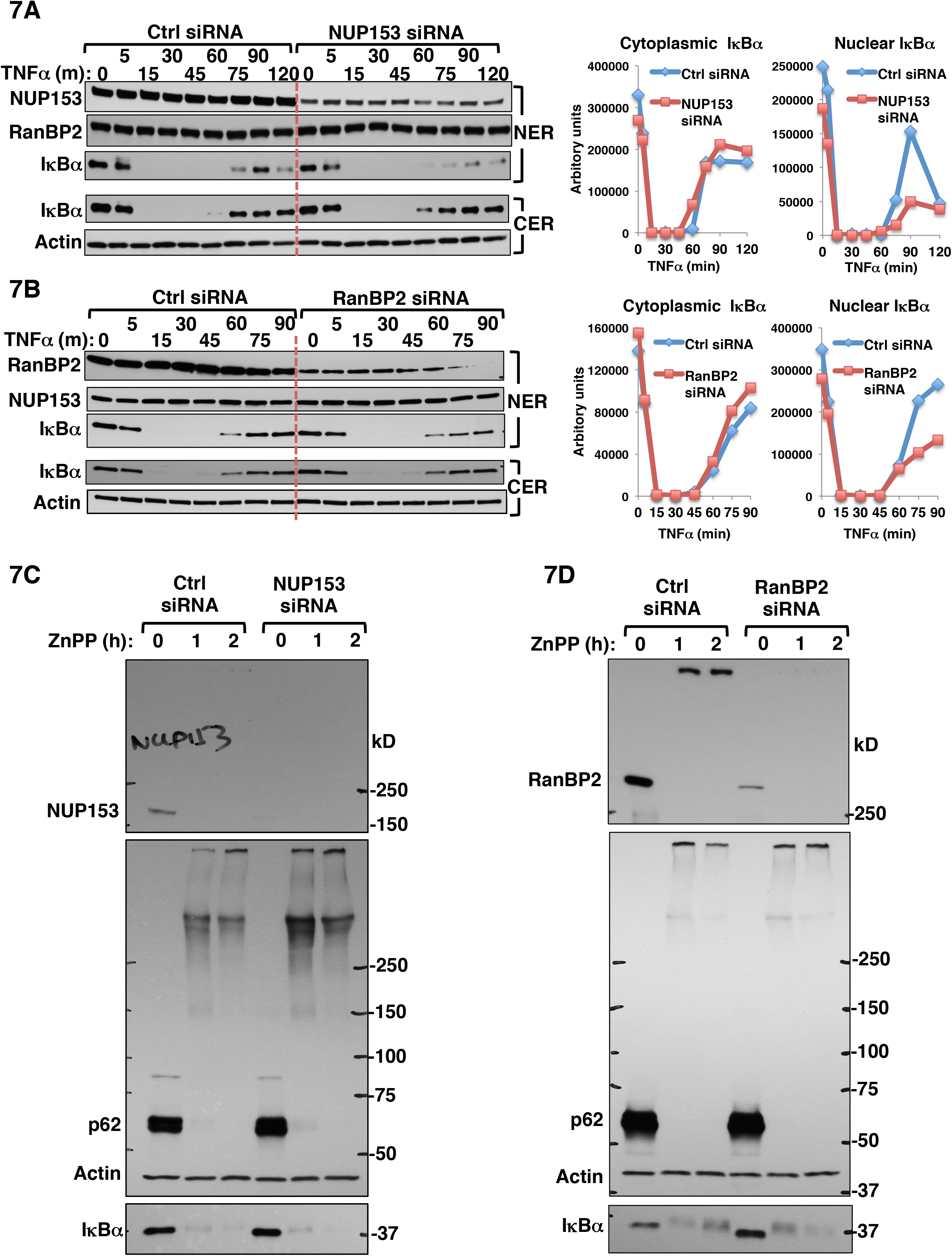
Effects of siRNA knockdown of Nup153 (A) and RanBP2 (B) on TNFα-elicited IκBα nuclear import. HEK293T cells were transfected with control siRNA (Ctrl) or NUP153 siRNA (**A**) or RanBP2 siRNA (**B**) or for 48 h, and then treated with 20 ng/ml TNFα for the indicated times. Cytosol and nuclear extracts were used for IB analyses. IκBα content was plotted over time. **C**. HEK293T cells were transfected with control siRNA (Ctrl) or NUP153 siRNA for 48 h, and then treated with 10 μM ZnPP for the indicated times. Cell lysates were used for IB analyses. **D**. HEK293T cells were transfected with control siRNA (Ctrl) or RanBP2 siRNA for 48 h, and then treated with 10 μM ZnPP for the indicated times. Cell lysates were used for IB analyses.

### IκBα siRNA knockdown analyses reveal that ZnPP-triggered NF-κB activation may additionally involve IκBβ-sequestration

Because of the functional redundancy of various IκBs in NF-κB cytoplasmic retention, IκBα-deficiency through genetic ablation, siRNA knockdown, or cycloheximide-inhibition of IκBα-synthesis fails to increase constitutive NF-κB transcriptional activation in many nonhepatic cell-types [19, 39, 40]. Thus, upon our IκBα siRNA-knockdown, a similar slight, barely detectable constitutive NF-κB activation was observed in HepG2 cells (Fig. 8A, left panel), whereas in primary mouse hepatocytes this activation was quite substantial (Fig. 8A, right panel). However, it was not quite as marked as that observed upon ZnPP-treatment, possibly due to the significant compensatory IκBβ-upregulation upon IκBα siRNA-knockdown (Fig. 8B). Because in hepatocytes both IκBα and IκBβ are the predominantly expressed IκBs [18, 21], and given the relatively more pronounced, sustained and consistently reproducible ZnPP-elicited NF-κB activation, we determined whether ZnPP also similarly sequestered IκBβ. Indeed, upon IκBα siRNA of HepG2 cells, the compensatorily increased IκBβ was similarly targeted to ZnPP-elicited aggregation, resulting in no appreciable mitigation of the ZnPP-induced NF-κB activation post IκBα-knockdown (Fig 8C). Furthermore, immunoblotting re-analyses of the RIPA-soluble ZnPP-treated lysates revealed that IκBβ also disappeared from its usual monomeric position, but was found in HMM protein aggregates (Fig. 8D). Such an apparent “IκBβ-loss” was also resistant to various UPD, calpain and ALD inhibitors, consistent with a sequestration process similar to that of IκBα. The ZnPP-targeting of both hepatic IκBs thus accounts for its potent NF-κB activation. Accordingly, *in vivo* ZnPP-treatment of mice revealed marked mRNA increases of several hepatic proinflammatory NF-κB target genes i.e. IL-6, IL-1β, TNFα and osteopontin (Fig. 8D)

**Fig. 8.**
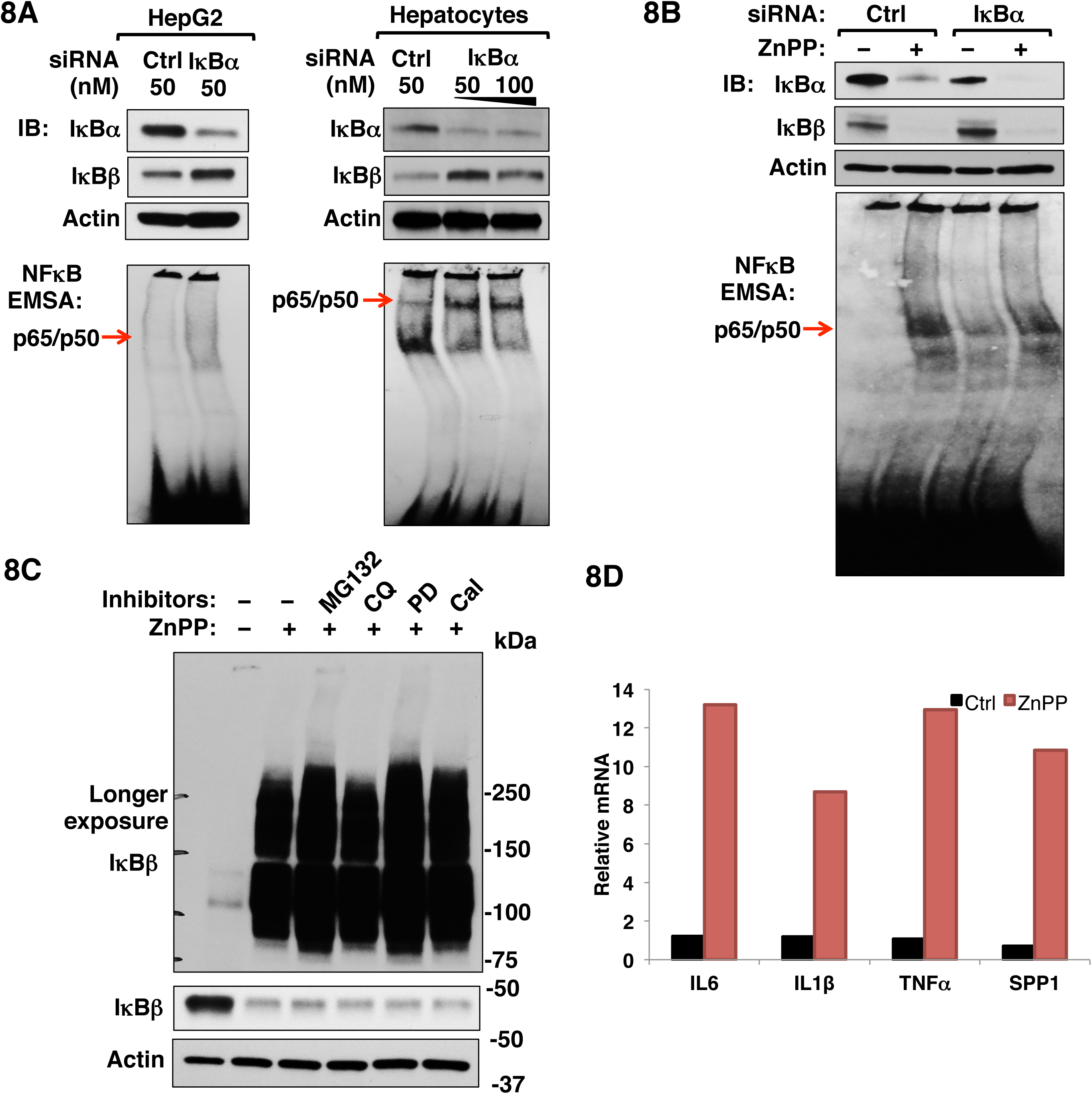
Effects of of IκBα siRNA knockdown on IκBβ-levels and constitutive (A) and ZnPP-induced NFκ B-activation (B). siRNA knockdown and EMSA analyses were conducted as detailed in Supplementary Methods. The marked hepatic IκBβ-upregulation coupled with the relatively less pronounced NFκB-activation observed upon IκBα-siRNA knockdown versus ZnPP-treatment led to the examination of whether IκBβ was similarly sequestered by ZnPP as IκBα (**C**). Proteolytic inhibitors used as in Fig. 2D. (**D**). qRT-PCR analyses of mRNA from intact livers of mice injected i.p., daily with ZnPP (50 μmol/kg) or vehicle controls (Ctrl) for 7 d (Mean, n = 2). SPP1, osteopontin or secreted phosphoprotein 1.

## DISCUSSION

### Why is IκBα vulnerable to ZnPP-elicited physical sequestration from the cytoplasm?

Recent evidence increasingly supports a sequestration and co-aggregation model of pathogenesis [41]. In a snowballing effect, aggregation-prone disease proteins “hijack” their interacting partners with vital functions, sequestering them into cytotoxic insoluble inclusions [41]. The protein scaffold p62 is one such IκBα-co-aggregating protein. However, scrutiny of ZnPP-treated p62^-/-^ hepatocytes and MEF cells reveals that it is not essential for ZnPP-elicited IκBα-sequestration. This is consistent with the report that in p62^-/-^ mouse liver, p62 is not required for MDB formation, just for their maturation and stability [42]. These aggregation-prone proteins share some common physicochemical properties: Preexistent proteins are relatively large in size, enriched in domains with high intrinsic disorder or unstructured regions, and exhibit low average hydrophobicity [43]; newly synthesized proteins on the other hand, are more vulnerable due to prolonged exposure of their relatively hydrophobic domains either during or soon after synthesis during their subsequent folding, assembly, or transport [43].

None of the 47 proteins that interacted with IκBα and co-aggregated upon ZnPP-treatment is a well-established stable IκBα interactor. Although most of the known stable IκBα interactors were repeatedly found in our co-IP analyses (Fig 4C), they were never detected in the ZnPP-aggregate proteome. This suggests that these 47 proteins may be transient IκBα interactors that are difficult to capture without highly sensitive and selective approaches. Our analyses of the physicochemical properties of these 47 proteins indicated that they ranged widely in size, with ∼90% exhibiting relatively low hydrophobicity, and ∼80% predicted to contain a long (>30 residues) disordered segment (Table S1). These dual features of low hydrophobicity coupled with high intrinsic disorder could synergistically contribute to their ZnPP-elicited co-aggregation upon transient interaction, and may explain why ZnPP initially targets preexistent IκBα species exhibiting both features (Table S1). The structurally related IκBβ is similarly vulnerable to ZnPP-elicited sequestration thus accounting for the remarkably profound NF-κB-activation. Whether this vulnerability stems from their common structural ankyrin-repeat domain feature remains to be determined. Free IκBα species is apparently unstable and requires NF-κB binding for folding to a stable conformation [44]. After TNFα stimulation, newly synthesized IκBα would be unstable and thus greatly susceptible to ZnPP-elicited aggregation without nuclear import in order to bind NF-κB and properly fold, thus resulting in the sustained ZnPP-triggered NF-κB-activation.

Although NF-κB and IκBα form a stable complex, they are not static cytoplasmic residents but shuttle continuously between the nucleus and cytoplasm [45]. Interestingly, p65 and IκBα translocate into the nucleus via different pathways. p65 enters the nucleus via the classical NLS-and importin (α3/α4)-dependent machinery [46], whereas IκBα and several other ankyrin repeat domain-containing proteins (ARPs) enter the nucleus via a NLS- and importin-*independent* machinery [47]. This machinery depends on a direct interaction between the ARPs, RanGDP as well as mobile nucleoporins such as Nup153 and RanBP2 for entry across the NPC into the nucleus. Indeed, plausibly relevant to this IκBα-nucleocytoplasmic shuttling process, our proteomic analyses of the 47 common IκBα-interacting and ZnPP-co-aggregating proteins underscored significant enrichment of mobile NPC nucleoporins. Of these, our heat-map analyses singled out Nup153 as both the top-ranked IκBα-interactant and ZnPP-aggregation target (Fig. 5B). By contrast, importins α3 and α4 relevant to p65 nuclear import were not found in the ZnPP-aggregate proteome, which may explain why p65 is not co-aggregated during its nuclear import.

### Potential role of Nup153 in IκBα-mediated NF-κB-repression

The nucleoporin Nup153 primarily exists N-terminally anchored to the nuclear pore basket while its disordered and flexible FG-rich C-terminus extends into the cytoplasm [48]. Another Nup153 pool apparently exists that shuttles between the cytoplasmic and NPC faces [49], and interacts directly with cargos (i.e. Stat1, Smad2, and PU.1) for their nuclear import via a transporter-*independent* machinery [48]. Nup153, in contrast to other FG-rich nucleoporins i.e. Nup98, Nup62, Nup214, also strongly binds ARs of iASPP and ASPP2, two representative substrates of the NLS-independent ARP-RanGDP nuclear import pathway [47]. Structural/biochemical analyses indicate that Nup153 binds RanGDP at a higher affinity than RanGTP [50]. This collective evidence suggests a critical role of Nup153 in facilitating ARP-RanGDP nuclear import. Our findings that IκBα co-immunoprecipitated not only with Nup153 in the nuclear extracts (NER), but also more appreciably with that in the cytoplasmic extracts (CER), even though the basal cytoplasmic Nup153-content is much lower than its nuclear content (compare inputs, Fig. 6B), are consistent with a minor, albeit highly dynamic, cytoplasmic Nup153 pool. This pool directly interacts with IκBα and may be responsible for the continuous IκBα-nucleocytoplasmic traffic at steady state, as indeed verified by our Nup153 knockdown analyses (Fig. 7). The concurrent ZnPP-elicited aggregation of Nup153 and IκBα suggests that the basal nuclear import of IκBα as well as that of newly synthesized IκBα after TNFα-stimulation is disrupted, leading to NF-κB nuclear persistence and prolonged response.

Another noteworthy proteomic finding is our quite prominent detection of the SUMO E3-ligase RanBP2 in this 47-protein cohort. RanBP2, an outer NPC component, fans its FG-rich filaments into the cytoplasm [51]. The proteomic detection of RanBP2, but no other cytoplasmic NPC nucleoporin i.e. Nup214 in this common 47-protein cohort is functionally intriguing because besides Nup153, RanBP2 is also involved in receptor-independent nuclear import through direct interactions with its specific cytoplasmic cargos [51]. This is intriguing given that IκBα SUMOylation is reportedly involved in its nuclear entry and subsequent post-induction NF-κB-repression [52]. Because RanBP2 also strongly binds RanGDP, conceivably a RanBP2-RanGDP-Nup153-IκBα-SUMO complex is involved in IκBα-nuclear import, a possibility consistent with our RanBP2 knockdown analyses (Fig. 7). The inviability of actively dividing HEK293T cells precluded our simultaneous knockdown of both nucleoporins to mimic their concurrent ZnPP-elicited sequestration. The failure of Nup153 or RanBP2 knockdown to affect ZnPP-elicited IκBα-sequestration suggests that the newly synthesized IκBα is itself vulnerable, particularly in the absence of both Nup153 and RanBP2 to chaperone its nuclear entry.

We find it quite instructive that many of these nucleocytoplasmic trafficking proteins as well as others within the 47 co-aggregated proteome are well-established components of pathogenic neurodegenerative disease protein aggregates (MS and ALS) or inclusions (Parkinson’s Lewy bodies), as well as hepatic MDBs (liver disease and hepatic protoporphyria models) [Fig. 5B; [13, 36, 37, 43, 53, 54]]. For example, Nup153 was shown not only to co-aggregate with cargo proteins in oligodendrocyte precursor cells in MS lesions [55], but also detected as anterior horn cell cytoplasmic inclusions of an ADAR2-deficient ALS mouse model [56]. These findings not only underscore the common intrinsic physicochemical and physiological properties of these aggregation-prone proteins, but also suggest that their interaction with IκBα could also similarly drive their co-aggregation under disease stresses, and thus account for the associated tissue inflammation and injury commonly seen in both liver and neurodegenerative diseases.

### Plausible clinical relevance to EPP and XLPP

Such ZnPP-elicited persistent NF-κB transcriptional activation may also account for the inflammation and injury stemming from chronic PPIX exposure not only in Fech^m1Pas^, DDC- and GF-protoporphyric rodent livers, but also in EPP and XLPP patient livers [10, 11, 28-30, 57], wherein avid Fe-overutilization may drive ZnPP-generation. ZnPP generation is also a prominent feature of iron-deficiency anemias and lead poisoning (Lamola and Yamane, 1974), Although our studies were largely confined to primary hepatocytes, in intact protoporphyric human and mouse livers these parenchymal cells are juxtaposed to non-parenchymal Kupffer cells and hepatic stellate cells that are quite capable of both canonical and non-canonical NF-κB-signaling. Thus, upon chronic PPIX-exposure, their NF-κB-activation could induce proinflammatory cytokines, chemokines, growth factors and other mitogens [17]. This additional paracrine cytokine stimulus could further potentiate the PPIX-elicited NF-κB-signaling activation within the hepatocyte, further aggravating the extent of EPP and XLPP liver injury. Indeed, exogenous TNFα greatly accelerated ZnPP-elicited IκBα-sequestration in hepatocytes (Fig. 6D)

The precise mechanism of PPIX-induced protein aggregation remains to be elucidated. Porphyrins trigger protein cross-linking through a secondary reaction between the photooxidative products of histidine, tyrosine and/or tryptophan and free NH_2_-groups of amino acids [58]. Given the light-sheltered *in vivo* environment of the liver, such a photooxidative protein crosslinking seems unlikely in PPIX-mediated hepatic MDB formation, although PPIX can trigger protein aggregation and cell toxicity in the dark, albeit at a slower rate [36]. Additionally, porphyrins also induce structural changes in certain proteins, leading to their functional impairment [59]. Such structural changes could expose some hydrophobic regions thereby inducing protein aggregation. In this context, the 192 PPIX-targeted aggregate proteins identified herein, we propose, could provide valuable insight into the future elucidation of the precise mechanism of PPIX-induced protein aggregation.

### Experimental Procedures

#### Cell culture and transfections

Primary mouse hepatocytes were isolated from C57BL/6 wild type male mice by collagenase perfusion. HepG2, HeLa, HEK293T and MEF cells were cultured using standard methods. HEK293T cells were transfected using TurboFect (ThermoFisher) and HepG2 cells were transfected using X-tremeGENE HP (Roche) according to the manufacturers’ instructions. See Extended Experimental Procedures for details.

#### EMSA

Nuclear fractions of cells were prepared using NE-PER Nuclear and Cytoplasmic Extraction Reagents (ThermoFisher). EMSA were performed using LightShift Chemiluminescent EMSA Kit (ThermoFisher). See Extended Experimental Procedures for details.

#### Western Immunoblotting (IB) Analyses

Whole-cell extracts were prepared with Cell Lysis buffer (Cell Signaling Technology) supplemented with 10% glycerol and protease/phosphatase inhibitor cocktail (Pierce). Cell lysates were sonicated for 10s and then cleared by centrifugation at 4°C in a tabletop centrifuge at the highest speed for 10 min. Protein concentrations were determined by BCA assay and equal amounts of proteins were separated on Tris-Glycine eXtended (TGX) polyacrylamide gels. Proteins were transferred onto nitrocellulose membranes (Biorad, Hercules, CA) for IB analyses. See Extended Experimental Procedures for details.

#### Mass spectrometry (MS)

Samples subject to MS proteomics analyses were processed using standard in-gel digestion and the resulting peptide mixture was desalted using C18-Ziptips (Millipore) and then injected into an LTQ-Orbitrap Velos mass spectrometer (Thermo Fisher). Raw data were processed using ProteinProspector (version 5.19.1; http://prospector.ucsf.edu/prospector/mshome.htm). See Extended Experimental Procedures for details.

## Acknowledgments

We gratefully acknowledge Mr. Chris Her, UCSF Liver Cell & Tissue Biology Core Facility (supported by NIDDK Grant P30DK26743) for hepatocyte isolation. We are most grateful to Dr. D. M. Bissell (UCSF) for valuable discussions and his critical review of our manuscript. We also gratefully acknowledge Prof. P. Ortiz de Montellano (UCSF) for valuable discussions of porphyrin chemistry, and Dr. G. Knudsen (UCSF) for her helpful comments on our IAP/MS methodology and critical review of our manuscript and proteomic data. We also sincerely thank Profs. T. Ishii, M. Komatsu, H. Zhu, N. Mizushima, R. Scheckman, and R. J. Youle for valuable cell-lines. Y. Liu is most grateful to Dr. D. Larsen, UCSF Nikon Imaging Center for her training in confocal immunofluorescence microscopy, and Drs. M. Laurance, UCSF HDF Comprehensive Cancer Center and A. Pico, The Gladstone Institute Bioinformatics Facility, for training on pathway analyses and visualization.

## Author Contributions

Y.L and M.A.C designed the studies and wrote the manuscript. M.A.C supervised the project. Y.L conducted most of the experiments with MS support and interpretation from M.T. and S.G. D.K. carried out the ZnPP *in vivo* mouse experiments. D.Y.K, J.J.C, and P.A.G provided critical, *sine qua non* research materials. All authors critically reviewed the manuscript.

## Supplementary Results & Discussion

### Results

#### NMPP-elicited IκBα-loss is independent of HRI eIF2α-kinase activation

The concurrence of IκBα loss and NF-κB activation and its attenuation by hemin-treatment led us to consider whether NMPP-elicited heme depletion was responsible for these findings. Conceivably, heme depletion could activate the hepatic heme-sensor HRI eIF2α kinase resulting in the translational suppression of IκBα and consequent NF-κB-activation. Similar IκBα translational suppression and consequent NF-κB-activation has been documented upon specific activation of the other cellular eIF2α kinases [1, 2]. Indeed, in parallel with IκBα-loss (Fig S2A), NMPP treatment increased the relative ratio of phosphorylated eIF2α (eIF2αP) over basal eIF2α levels, indicating HRI activation [3]. However, NMPP-treatment of cultured hepatocytes from cultured hepatocytes from HRI WT (HRI^+/+^) and knockout (KO; HRI^-/-^) mice elicited a comparable IκBα, excluding causal HRI activation (Fig. 1D).

#### ZnPP-elicited IκBα-loss is independent of autophagy, calpain-mediated proteolysis and reactive oxygen species (ROS)

Given the reported IκBα-degradation via ALD [4] and the known p62-role in this process [5], we conclusively excluded ALD in this process by documenting that knockout of the essential autophagic gene ATG5 failed to mitigate ZnPP-elicited IκBα loss (Fig. S2B). Because of possible calpain-mediated IκBα-degradation [6, 7], we also excluded its involvement by documenting that knockout of the calpain-degradation pathway (capn4 KO MEF cells) failed to abrogate ZnPP-elicited IκBα loss (Fig. S2C). These findings coupled with those with ALD and calpain inhibitors (Fig. 2D) revealed that neither ALD nor calpain degradation played any role in this IκBα loss. Furthermore, through the use of various ROS-quenchers (Fig. S3), we excluded any possible PPIX-photoactivation and consequent ROS-mediated oxidative stress in this IκBα-loss [8].

#### Discussion

The precise mechanism of the ZnPP-elicited IκB-protein sequestration is currently unknown, but some plausible mechanisms are presented. The fact that newly synthesized IκBα is particularly vulnerable to such ZnPP-sequestration implicates the involvement of its intrinsic disordered domains at a stage when it has not yet reached its mature folded state.

In search of clues on potential ZnPP-mechanisms of protein sequestration, we sought the advice of various internationally recognized expert heme/porphyrin chemists and biochemists and porphyria experts on the possible mechanistic causes. Not a single investigator was aware that ZnPP caused protein aggregation, much less IκBα-sequestration. ZnPP (along with SnPP), is actually believed to be safe and is recommended for the treatment of neonatal jaundice due to its effective inhibition of microsomal heme oxygenase (HO1), the key rate-limiting enzyme in the conversion of heme to biliverdin [9]. However, *in vivo*, overproduction of protoporphyrin IX (PPIX) due to defects in ferrochelatase (congenital erythropoietic protoporphyria) or PPIX-overproduction in X-linked protoporphyria, as well as iron-deficiency or lead-poisoning induced anemias (that can exhaust iron-stores) leads to ZnPP-generation [10-12]. Our findings suggest that under these conditions, these patients may easily succumb to ZnPP-elicited IκBα-sequestration and consequently unabated hepatic NF-κB-elicited activation of cytokines and chemokines. We believe, these findings would for the first-time alert physicians of this pathological potential of ZnPP and are clinically relevant not just in MDB-inducing diseases but also in clinical protoporphyrias and iron-deficiency/lead-induced anemias.

## Supplementary methods

### Contact for reagent and resource sharing

Information and requests for resources and reagents should be directed to M. A. Correia.

### Experimental model details

#### Animal studies

C57BL/6 wild type male mice (8-12-week old) purchased from the Jackson Laboratory (Bar Harbor, ME). Mice were fed a standard chow-diet and maintained under a normal diurnal light cycle. All animal experiments were carried out strictly by protocols specifically approved by the UCSF/Institutional Animal Care and Use Committee (IACUC) and its care and use of laboratory animal guidelines. For the ZnPP-treatment, ZnPP was dissolved in sterile DMSO at 100 mM, and then diluted 1:10 (v:v) with sterile 7.5% BSA to a final concentration of 10 mM. Mice were weighed and then injected daily with either ZnPP (50 μmol/kg, i.p.), or vehicle (controls) (10% DMSO in 7.5% BSA) for 7 days. Liver samples were collected and frozen at −80°C until RNA isolation.

#### Cell culture

All cells were grown at 37°C with 5% CO_2_ in a humidified incubator. C57BL/6 wild type male mice (8-12-week old) purchased from the Jackson Laboratory (Bar Harbor, ME) were used for primary hepatocyte preparation. Hepatocytes were isolated by *in situ* collagenase perfusion and purified by Percoll-gradient centrifugation by the UCSF Liver Center Cell Biology Core, as described previously [13]. Fresh primary mouse hepatocytes were cultured on Type I collagen-coated 60 mm Permanox plates (Thermo Scientific, Grand Island, NY) in William’s E Medium supplemented with 2 mM L-glutamine, insulin-transferrin-selenium, 0.1% bovine albumin Fraction V, Penicillin-Streptomycin and 0.1 μM dexamethasone. Cells were allowed to attach for 4 to 6 h and then overlaid with Matrigel. From the 2nd day after plating, the medium was replaced daily, and cells were further cultured for 4-5 days with daily light microscopic examination for any signs of cell death and/or cytotoxicity. On day 5, some cells were treated with 30 μM NMPP (dissolved in DMSO), 10 μM PPIX (dissolved in DMSO with sonication) or 10 μM zinc (II)-PPIX (ZnPP, dissolved in DMSO and complexed with bovine serum albumin (BSA) at a molar ratio of 4:1 to keep it solubilized in the medium) for various times as indicated (Results). In some cases, hepatocytes were pretreated with various inhibitors as indicated (Results) for 1 h before treatment with NMPP, PPIX or ZnPP.

HepG2 and HeLa cells were cultured in minimal Eagle’s medium (MEM) containing 10% v/v fetal bovine serum (FBS) and supplemented with nonessential amino acids and 1 mM sodium pyruvate. HEK293T and MEF cells were cultured with Dulbecco’s Modified Eagle high glucose medium (DMEM) containing 10% v/v FBS. For transfection experiments, cells were seeded onto 6-well plates, and when cells were 60% confluent, each cell well was transfected with 3 μg plasmid DNA complexed with TurboFect transfection reagent for HEK293T cells and X-tremeGENE HP transfection reagent for HepG2 cells according to the manufacturers’ instructions. At 40-72 h after transfection, cells were either treated as indicated or directly harvested for assays.

### Method details

#### Hepatic PPIX content

PPIX content was determined using the intrinsic PPIX fluorescence as described [14]. Briefly, 50 μl of cell lysates were first extracted with 400 μl of EtOAc-HAc (3:1) and then re-extracted with another 400 μl of EtOAc-HAc. The extracts were pooled and reextracted with 400 μl of 3 M HCl. After centrifugation, the aqueous phase was recovered for fluorescent determination in a SpectrumMax M5 plate reader at an excitation 405 nm. The intensity at emission 610 nm was quantified relative to a standard curve prepared with known concentrations of pure PPIX.

#### Plasmids

The primers, templates, vectors and restriction enzymes (RE) used for constructing plasmids used in this study are summarized below:

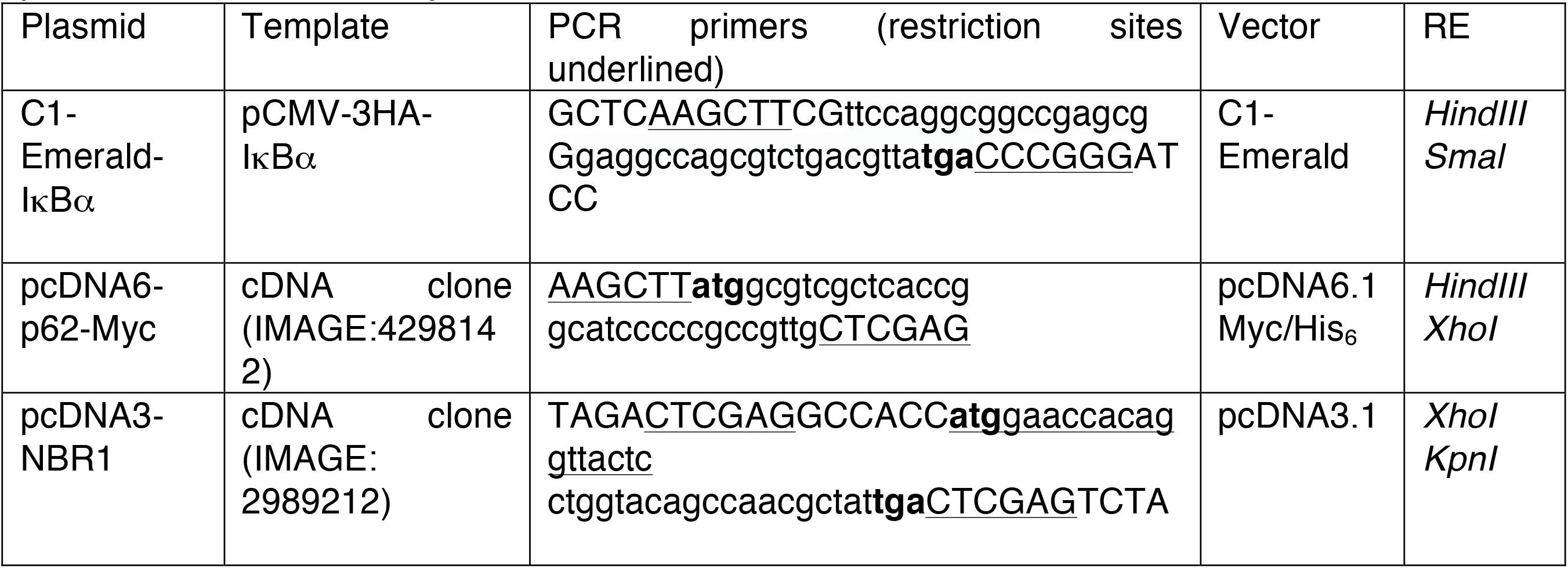

#### Electrophoretic mobility shift assay (EMSA)

Nuclear fractions of cells were prepared using NE-PER Nuclear and Cytoplasmic Extraction Reagents. EMSA were performed using LightShift Chemiluminescent EMSA Kit. Briefly, 2-5 μg of nuclear extracts were incubated with 1 μl of biotin-labeled NFkB DNA probes in binding buffer [10 mM Tris, pH 7.5, 50 mM KCl, 1 mM DTT, 5 mM MgCl_2_, 0.2 mM EDTA, 5% glycerol, 0.5% NP-40, 1 μg of poly dI-dC]. The final volume of the mixture was adjusted to 20 μl and incubated at room temperature for 30 min, and then mixed with 5 μl of loading buffer for loading onto 5% TBE Polyacrylamide Gel (Biorad, Hercules, CA). The gel was run at 120 V until the dye reached the gel bottom, and then transferred to Hybond-N+ positively charged nylon membrane (GE Life Sciences, Marlborough, MA). The NF-κB complex shifted probes were detected by blotting with HRP-coupled streptavidin. For super-shift EMSA, the binding mixture (in the absence of the biotin-labeled probes) was first incubated with p65 antibody for 20 min on ice, and then the biotin-labeled probes were added and further incubated at room temperature for another 30 min before gel loading.

#### RNA isolation and semi-quantitative RT-PCR

Total RNA was extracted using RNeasy Mini Kit according to the manufacturers’ instructions. Total RNA (2 μg) was used to perform reverse transcription using SuperScript VILO Master Mix in a 20 μl-reaction. Reverse transcribed first strand cDNA (1 μl) was used in PCRs.

#### Western Immunoblotting (IB) Analyses

For Western IB analysis, whole-cell extracts were prepared with Cell Lysis buffer containing 20 mM Tris-HCl (pH 7.5), 150 mM NaCl, 1 mM EDTA, 1 mM EGTA, 1% Triton, 2.5 mM sodium pyrophosphate, 1 mM β-glycerophosphate, 1 mM Na3VO4, 1 µg/ml leupeptin and supplemented with 10% glycerol and protease/phosphatase inhibitor cocktail. Cell lysates were sonicated for 10s and then cleared by centrifugation at 4°C in a tabletop centrifuge at the highest speed for 10 min. Protein concentrations were determined by BCA assay and equal amounts of proteins were separated on 4-15% Tris-Glycine eXtended (TGX) polyacrylamide gels. Proteins were transferred onto nitrocellulose membranes (Biorad, Hercules, CA) for IB analyses. The antibodies used are listed in Key Resources Table.

#### Co-Immunoprecipitation (Co-IP) analyses

Whole-cell extracts were prepared as described above. Cell lysates (1 mg) were then incubated with indicated antibodies (2 µg) or control IgGs at 4°C overnight. Antibody-antigen complexes were then captured by protein G Dynabeads at room temperature for 1 h, and then eluted by heating at 95°C for 10 min in 2X SDS-loading buffer. Eluates were subjected to IB analyses as described above.

#### Sequential solvent extraction of cell lysates

Cells were harvested in cell lysis buffer as described above and the cell lysates were cleared by centrifugation at 14,000g. The pellet was then solubilized in RIPA buffer supplemented with 0.1% SDS, 10% glycerol and protease/phosphatase inhibitor cocktail with sonication followed by centrifugation at the highest speed for 10 min. The resulting pellet was then solubilized with sonication in urea/CHAPS buffer containing 8 M urea, 2 M thiourea, 4% CHAPS, 20 mM Tris-base, and 30 mM DTT and supplemented with protease/phosphatase inhibitor cocktail. High salt buffer (HSB)-extraction (Fig. 4C) was carried out as described previously [15]. Briefly, cells were first harvested in cell lysis buffer containing 1% Triton as described above, the resulting pellet was then suspended in a cell lysis buffer supplemented with 1.5 M KCl (HSB) with sonication. Upon sedimentation, the resulting pellet was solubilized in Laemmli buffer containing 4% SDS or in urea/CHAPS buffer as described above.

#### Confocal Immunofluorescence microscopy (CIFM)

Cells were grown on collagen-coated glass coverslips and treated as indicated. Cells were fixed with 4% formaldehyde for 20 min at room temperature followed by methanol at −20°C for 1 min. After that, cells were rinsed with PBS and blocked for 1 h with 10% normal goat serum in PBS /0.1% Tween at room temperature, and then stained with indicated primary antibodies at 4°C overnight. Cells were then washed in PBS 0.1% Tween three times and then stained with secondary antibodies for 1 h at room temperature. Cells were further washed three times in PBS 0.1% Tween and then mounted using ProLong Diamond Antifade Mountant with DAPI nuclear stain (Molecular Probes, Grand Island, NY). The following secondary antibodies were applied: Goat anti-rabbit IgG Alexa Fluor 488 (Invitrogen, Grand Island, NY), anti-mouse IgG Alexa Fluor 647 (Cell Signaling Technology, Danvers, MA). These particular fluor dyes were selected to circumvent any specific interference from ZnPP intrinsic fluorescence. Images were taken with a Nikon Yokogawa CSU-22 Spinning Disk Confocal Microscope using a Plan Apo VC 100x/1.4 oil lens or on a Nikon high-speed wide-field Andor Borealis CSU-W1 spinning disk confocal microscope using a Plan Apo VC 60x/1.4 oil lens. Images were processed using ImageJ software. For quantification, at least 600 cells at each condition were evaluated. Statistical significance was tested using two-sided unpaired Student’s t-test.

#### Immunoaffinity purification (IAP)

We employed high-affinity alpaca Nanobody crosslinked beads (GFP-Trap) for IAP, which not only enabled a high-level enrichment of target proteins but also eliminated IgG contaminants that confound downstream LC-MS/MS analyses. N-terminally mEmerald (GFP)-tagged IκBα (GFP-IκBα) was transiently transfected into HEK293T cells, with mEmerald-transfected cells as background control (C1-GFP). Two wells of HEK293T cells grown on 6-well plates were pooled and lysed in 1 ml lysis buffer supplemented with 10% glycerol, protease/phosphatase inhibitor cocktail and 20 mM N-ethylmaleimide (NEM). Centrifugation-cleared cell lysates were incubated with 50 μl GFP-trap agarose beads at 4°C overnight. Subsequently, GFP-trap beads were collected by centrifugation at 3,000g for 30s and then washed 5 times with cell lysis buffer. Co-immunoprecipitated proteins were eluted by incubating beads with 2X Laemmli buffer at 70°C for 15 min. Eluates were then subjected to SDS-PAGE and stained with Coomassie Blue to visualize the bands for subsequent in-gel digestion.

#### Biotinylation by antibody-recognition (BAR)

Traditional approaches for in vivo protein-protein interactions such as co-IP are based on affinity capture of stable protein complexes that will be disrupted under harsh denaturing conditions. By contrast, biotinylation proximity labeling circumvents this limitation by introducing an enzyme to the target protein that can generate distance-constrained reactive biotin molecules to covalently link neighboring proteins, providing a permanent tag that survives purification under harsh conditions for downstream identification [16]. We chose BAR/APEX over BioID (the two popular methods for in vivo biotinylation proximity) because the BAR approach [17] is much more robust as it enables quicker capture of interacting proteins (as the peroxidase mediated biotinylation takes only a few minutes) without the more protracted BioID methodology (requiring 18-24 h tagging time), which could overlook short-term/transient interactants. BAR analyses were performed according to [17]. Briefly, two wells of HEK293T cells grown on collagen-coated 6-well plates were first treated as indicated and then fixed with 4% formaldehyde for 10 min at room temperature and permeabilized for 7 min with 0.5% Triton-X in PBS. After rinsing with PBS, cells were incubated with 0.5% H_2_O_2_ for 10 min. After rinsing with PBS, cells were then blocked for 1 h with 10% normal goat serum in PBS/0.1% Tween, and then stained with IκBα antibody (mouse monoclonal, 44D4) at a 1:100 dilution in 1% normal goat serum in PBS/0.1% Tween at 4°C overnight. Negative control staining with no antibody was also performed. Cells were then washed in PBS 0.1% Tween for over 1 h with at least 5 buffer changes and then stained with secondary poly-HRP-conjugated goat anti-mouse IgGs (from Biotin XX Tyramide SuperBoost™ Kit) at a 1:1000 v:v dilution in 1% normal goat serum in PBS/0.1% Tween for 1 h. Cells were further washed for over 2 h in PBS 0.1% Tween with at least 5 buffer changes. After that, cells were labeled with biotin using Biotin XX Tyramide SuperBoost™ Kit following product instructions. Briefly, cells were pre-incubated with 300 ml of reaction buffer containing biotin-XX-tyramide (Biotin-Phenol) for 15 min; H_2_O_2_ was then added to obtain a final concentration of 0.5 mM and incubated for another 5 min. Negative controls without H_2_O_2_ were also included. The labeling reaction was then stopped by quickly exchanging the reaction solution with 1 ml of 500mM sodium ascorbate. Cells were washed 3 times with PBS and then lysed in RIPA buffer supplemented with 2% SDS and boiled for 1 h to reverse formaldehyde crosslinking. The cell lysates were cleared by centrifugation at 14,000g, the supernatants were used as whole cell lysates (WCL). The resulting pellet was further solubilized in urea/CHAPS buffer as described above. Aliquots from both fractions were used for streptavidin-HRP blot (Invitrogen, Grand Island, NY, #S911), to characterize the specificity and efficiency of BAR. The remaining WCL and urea/CHAPS fractions were separately diluted 1:2 (v;v) in RIPA buffer for streptavidin (SA) pull-down. SA pull-downs were performed according to [18]. Briefly, 50 μl of streptavidin magnetic beads were added to the lysates to incubate at 4°C overnight on a rotator. Beads were then washed twice with RIPA buffer, once with 1 M KCl, once with 2 M urea in 20 mM Tris base, and lastly, twice with RIPA buffer. Biotinylated proteins were then eluted by incubating beads in 40 μl of 2X Laemmli buffer supplemented with 2 mM biotin at 95°C for 5 min. Eluates were then subjected to SDS-PAGE and stained with Coomassie Blue to visualize the bands for subsequent in-gel digestion. For SA pull-down from urea/CHAPS fractions, the last 2 washes were performed using 100 mM ammonium bicarbonate (ABC) before proceeding with on-beads digestion.

#### Mass spectrometry (MS)

For in-gel digestion, each gel lane was sliced into 8-10 sections and processed separately using a standard in-gel digestion procedure [19, 20]. Briefly, each gel piece was reduced and alkylated, and then digested overnight with 300 ng of Trypsin/Lys-C Mix (Promega, Madison, WI). For on-bead digestion after SA pull-down, beads were first resuspended in 6 M urea in 100 mM ABC, and reduced by adding final 10 mM DTT and incubation at 37°C for 30 min. The samples were then alkylated by adding final 15 mM iodoacetamide with incubation at room temperature in the dark for 30 min. Trypsin/Lys-C Mix (100 ng) was then added and the mixture incubated for 4 h at 37°C. The mixture was then diluted 1:6 (v:v) with 100 mM ABC to reduce the urea concentration to 1 M and then further incubated at 37°C overnight. The resulting peptide mixture was desalted using C18-Zip tips (Millipore, Hayward, CA), speed-vacuumed to dryness and suspended in 0.1% formic acid for injection into an LTQ-Orbitrap Velos mass spectrometer (Thermo Fisher, Grand Island, NY) coupled to a nano-Acquity UPLC (Waters, Milford, MA) with an EASY-Spray column (75 µm x 15 cm column packed with 3 µm, 100 Å PepMap C18 resin; Thermo Scientific, Grand Island, NY). Each sample was separated on the column following a 90-min solvent gradient (Solvent A: Water/0.1% formic acid; Solvent B: Acetonitrile/0.1% formic acid) and the mass spectrometer was operated under high-energy collisional dissociation (HCD) mode. Peak lists were extracted using PAVA, an in-house software developed by UCSF Mass Spectrometry facility. Peak lists from 8-10 mass spectometric fractions under each experimental condition were pooled to search against the Swissprot human database (SwissProt.2015.12.1; 20194/549832 entries searched) using ProteinProspector (version 5.19.1; http://prospector.ucsf.edu/prospector/mshome.htm) [21]. A fully randomized database was used to estimate false discovery rates (FDR) [22]. Score thresholds were chosen at <1% FDR at the peptide level. ProteinProspector search parameters were as follows: Tolerance for precursor and product ions were 20 ppm and 25 ppm; a maximum of 1 missed cleavage of trypsin was allowed; carbamidomethylation of cysteine was set as a fixed modification; variable modifications include: N-terminal Met loss and/or acetylation, Met oxidation, peptide N-terminal Gln to pyroGly conversion; number of variable modifications was 2. Reporting thresholds were as follows: Minimum score of protein: 22.0; minimum score of peptide: 15.0; maximum E value of protein: 0.01; maximum E value of peptide: 0.05.

#### Mass spectral data processing

To obtain a high-confidence IkBa-interactome, three independent IAP-MS experiments were performed,(from cell transfection, IAP to MS protein identification),each consisting of three biological replicates (three wells each individually transfected). The identified proteins were strictly filtered as follows: 1) From each LC-MS/MS run, only proteins with two or more peptides identified were utilized for further analyses; 2) when some unique peptides matched to multiple protein isoforms, only the isoform with the highest number of matched peptides was reported; 3) only proteins that were immunoaffinity purified with mEmerald-IκBα in all three replicates were kept for further analyses. After filtering, an enrichment ratio of total spectral counts for each protein in mEmerald-IκBα IAP over GFP (C1-mEmerald) background control was calculated as described [23]. Proteins that were not detected in the background control were artificially assigned 0.5 spectral counts, so that an enrichment fold could be calculated. Only proteins that were enriched ≥ 2-fold in all three replicates were considered high-confidence interactors. Mean ratio and p-value from three replicates were then obtained. p-values were obtained using a paired Student’s t-test with total spectral counts from GFP-control compared to mEmerald-IκBα IAP values. To characterize the aggregate-proteome, proteins in high molecular mass (HMM)-regions and urea/CHAPS-solubilized fractions from HEK293T and HepG2 cells were similarly filtered and those in ZnPP-treated and control samples were directly compared. Identified proteins were filtered as described above. Overlap analyses were performed using the Venn Diagram tool (http://bioinformatics.psb.ugent.be/webtools/Venn/).

#### Network analysis of aggregated proteins

192 commonly found proteins in ZnPP-induced aggregates from both cell lines were searched against a STRING database version 10.0 [24] for protein-protein interactions. We used the highest confidence score (≥ 0.9) to obtain a protein-interaction network. The network was exported to Cytoscape software [25] for pathway annotation and visualization.

#### siRNA-knockdown analyses

HEK293T cells were transfected using DharmaFECT 4 Transfection Reagent, with 10 nM siRNA specific for RanBP2 or NUP153. Treatments were performed 48 h after transfection. HepG2 cells were transfected using DharmaFECT 4 Transfection Reagent with 50 nM siGENOME Human NFKBIA siRNA-Smartpool or control non-targeting siRNA. Primary mouse hepatocytes, were transfected using DharmaFECT 4 Transfection Reagent with 50-100 nM siGENOME Nfkbia siRNA-SMARTpool or control siRNA non-targeting siRNA, 16 h before overlay with Matrigel. Cells were harvested 72-96 h after transfection.

#### Bioinformatic Analyses

GRAVY scores estimating average hydrophobicity was calculated using Kyte and Doolittle hydropathic analyses (http://web.expasy.org/protparam/) [26]. Positive GRAVY scores suggest more hydrophobicity. Proteins with long intrinsically disordered regions was predicted using SLIDER method (http://biomine.cs.vcu.edu/servers/SLIDER/) [27]. This computational method is based on the physicochemical properties of amino acids, sequence complexity, and amino acid composition, a SLIDER score >0.538 suggests a high likehood that the protein contains a long (>30 residues) disordered segment.

#### Quantification and statistical analyses

Statistical significance was tested using the two-sided unpaired Student’s t-test. N, number of individual experiments, was indicated in each figure legend.

**Table 1.** Structural properties of the 47 IκBα**-**interacting and ZnPP-inducible-aggregating proteins.

| Gene | Accession | GRAVY | SLIDER | Protein Name |
| --- | --- | --- | --- | --- |
| NUP153 | P49790 | -0.469 | 0.869 | Nuclear pore complex protein Nup153 |
| NUP155 | O75694 | -0.129 | 0.701 | Nuclear pore complex protein Nup155 |
| IMMT | Q16891 | -0.465 | 0.835 | MICOS complex subunit MIC60 |
| EPRS | P07814 | -0.517 | 0.825 | Bifunctional glutamate/proline--tRNA ligase |
| EIF3A | Q14152 | -1.489 | 0.946 | Eukaryotic translation initiation factor 3 subunit A |
| LRPPRC | P42704 | -0.206 | 0.723 | Leucine-rich PPR motif-containing protein, mitochondrial |
| LBR | Q14739 | -0.056 | 0.651 | Lamin-B receptor |
| DDOST | P39656 | -0.057 | 0.417 | Dolichyl-diphosphooligosaccharide--protein glycosyltransferase 48 kDa subunit |
| RANBP2 | P49792 | -0.596 | 0.840 | E3 SUMO-protein ligase RanBP2 |
| ABCD3 | P28288 | -0.13 | 0.518 | ATP-binding cassette sub-family D member 3 |
| PDCD6IP | Q8WUM4 | -0.468 | 0.785 | Programmed cell death 6-interacting protein |
| MDN1 | Q9NU22 | -0.371 | 0.898 | Midasin |
| RPS2 | P15880 | -0.233 | 0.562 | 40S ribosomal protein S2 |
| MSH6 | P52701 | -0.499 | 0.836 | DNA mismatch repair protein Msh6 |
| KIF5B | P33176 | -0.81 | 0.876 | Kinesin-1 heavy chain |
| COPB1 | P53618 | -0.091 | 0.684 | Coatamer subunit beta |
| USP7 | Q93009 | -0.643 | 0.722 | Ubiquitin carboxyl-terminal hydrolase 7 |
| RPL10 | P27635 | -0.553 | 0.458 | 60S ribosomal protein L10 |
| HADHA | P40939 | -0.078 | 0.595 | Trifunctional enzyme subunit alpha, mitochondrial |
| DDX6 | P26196 | -0.323 | 0.674 | Probable ATP-dependent RNA helicase DDX6 |
| TRIM28 | Q13263 | -0.387 | 0.857 | Transcription intermediary factor 1-beta |
| COPG1 | Q9Y678 | -0.145 | 0.658 | Coatamer subunit gamma-1 |
| RPL8 | P62917 | -0.528 | 0.539 | 60S ribosomal protein L8 |
| HACD3 | Q9P035 | 0.003 | 0.390 | Very-long-chain (3R)-3-hydroxyacyl-CoA dehydratase 3 |
| NUP93 | Q8N1F7 | -0.366 | 0.705 | Nuclear pore complex protein Nup93 |
| HADHB | P55084 | -0.072 | 0.528 | Trifunctional enzyme subunit beta, mitochondrial |
| MCM3 | P25205 | -0.564 | 0.879 | DNA replication licensing factor MCM3 |
| POLR1C | O15160 | -0.28 | 0.416 | DNA-directed RNA polymerases I and III subunit RPAC1 |
| CAND1 | Q86VP6 | -0.019 | 0.750 | Cullin-associated NEDD8-dissociated protein 1 |
| EIF3M | Q7L2H7 | -0.184 | 0.544 | Eukaryotic translation initiation factor 3 subunit M |
| SLC25A11 | Q02978 | 0.117 | 0.301 | Mitochondrial 2-oxoglutarate/malate carrier protein |
| BIRC6 | Q9NR09 | -0.138 | 0.870 | Baculoviral IAP repeat-containing protein 6 |
| PSMA4 | P25789 | -0.459 | 0.609 | Proteasome subunit alpha type-4 |
| GTF2I | P78347 | -0.526 | 0.798 | General transcription factor II-I |
| IK | Q13123 | -1.427 | 0.928 | Protein Red |
| SLC25A10 | Q9UBX3 | 0.148 | 0.327 | Mitochondrial dicarboxylate carrier |
| KPNB1 | Q14974 | -0.092 | 0.704 | Importin subunit beta-1 |
| RPS6 | P62753 | -0.945 | 0.820 | 40S ribosomal protein S6 |
| RPL7A | P62424 | -0.547 | 0.685 | 60S ribosomal protein L7a |
| PSMD3 | O43242 | -0.529 | 0.821 | 26S proteasome non-ATPase regulatory subunit 3 |
| SEC16A | O15027 | -0.607 | 0.924 | Protein transport protein Sec16A |
| SEC23A | Q15436 | -0.238 | 0.552 | Protein transport protein Sec23A |
| RPS8 | P62241 | -1.02 | 0.651 | 40S ribosomal protein S8 |
| MCM5 | P33992 | -0.352 | 0.671 | DNA replication licensing factor MCM5 |
| GTF3C3 | Q9Y5Q9 | -0.363 | 0.858 | General transcription factor 3C polypeptide 3 |
| IPO4 | Q8TEX9 | 0.07 | 0.797 | Importin-4 |
| XRN2 | Q9H0D6 | -0.651 | 0.790 | 5'-3' exoribonuclease 2 |
| NFKBIA | P25963 | -0.452 | 0.8195 | NF-kappa-B inhibitor alpha |
| NFKBIB | Q15653 | -0.388 | 0.7360 | NF-kappa-B inhibitor beta |
| NFKBIE | O00221 | -0.4336 | 0.8077 | NF-kappa-B inhibitor epsilon |
| SQSTM1 | Q13501 | -0.632 | 0.6246 | Sequestosome-1 |

\* The GRAVY number of a protein is a measure of its hydrophobicity or hydrophilicity. The hydrophathy values range from -2 to +2 for most proteins, with the positively rated proteins being more hydrophobic. Less hydrophobic proteins are labeled red. \*\* Proteins are listed with SLIDER score, the higher the score the more likely a protein has a long ( $\geq 30$ AAs) disordered segment. Scores above 0.538 indicates that a given protein has a long disorder segment (labeled red).

---

**FIGURE S1.**
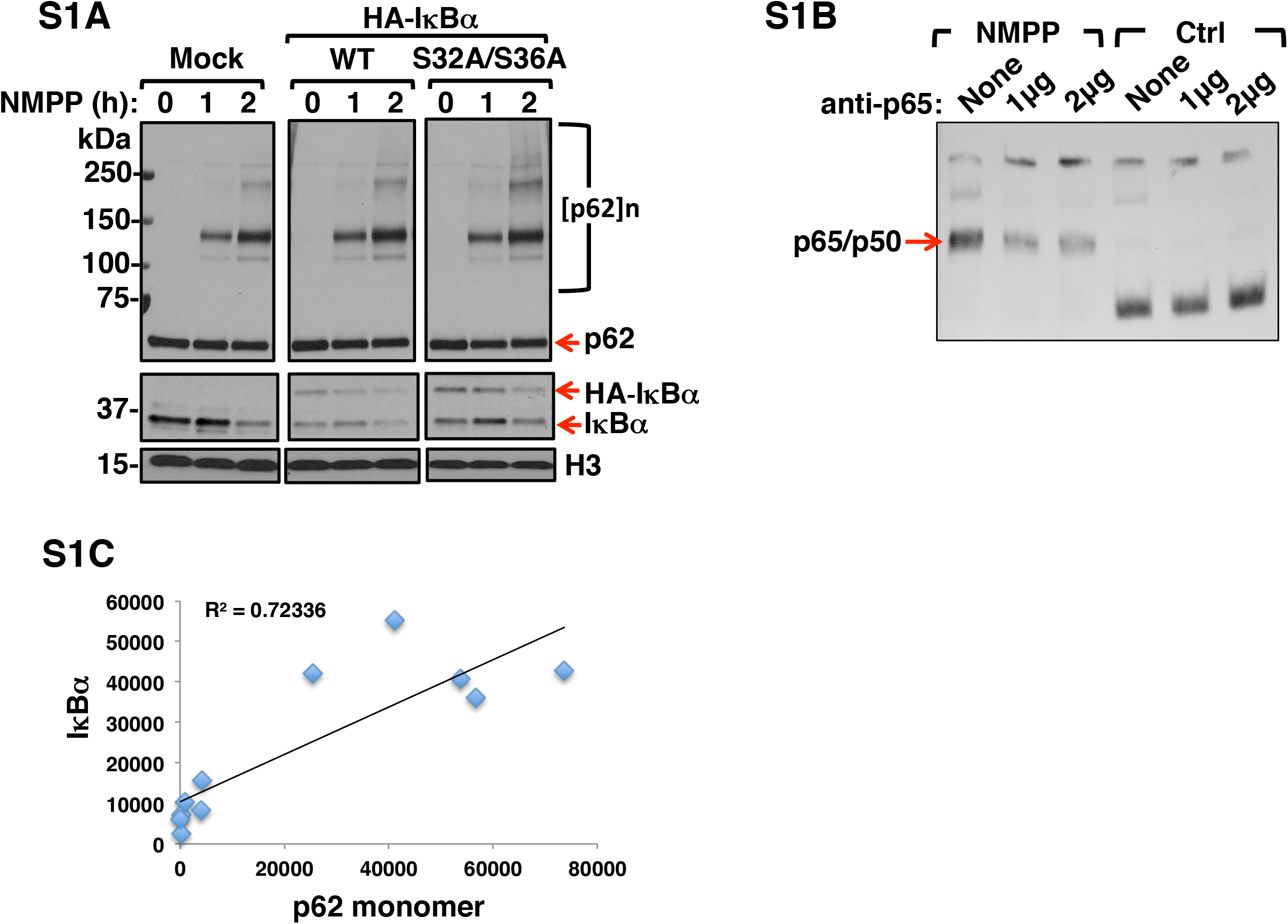
NMPP-elicited PPIX-accumulation with concurrent NFκB activation and IκBα-loss is independent of proteasomal degradation, PPIX and ZnPP are even more potent elicitors of IκBα-loss and p62-aggregation than NMPP (Related to Fig 1). **A.** HepG2 cells were transfected with pCMV4-3HA-IκBα or mutant pCMV4-3HA-IκBα-S32A/S36A for 40 h, then treated with NMPP for the indicated times. Cell lysates were subjected to IB analyses of p62 and IκBα with histone H3 as the loading control. **B**. Supershift EMSA using nuclear extracts from Fig 1D with p65-antibody verifies the identity of the p65/p50 band. **C**. The correlation of IκBα and p62 monomer levels quantified from Fig1E (n = 2).

**FIGURE S2.**
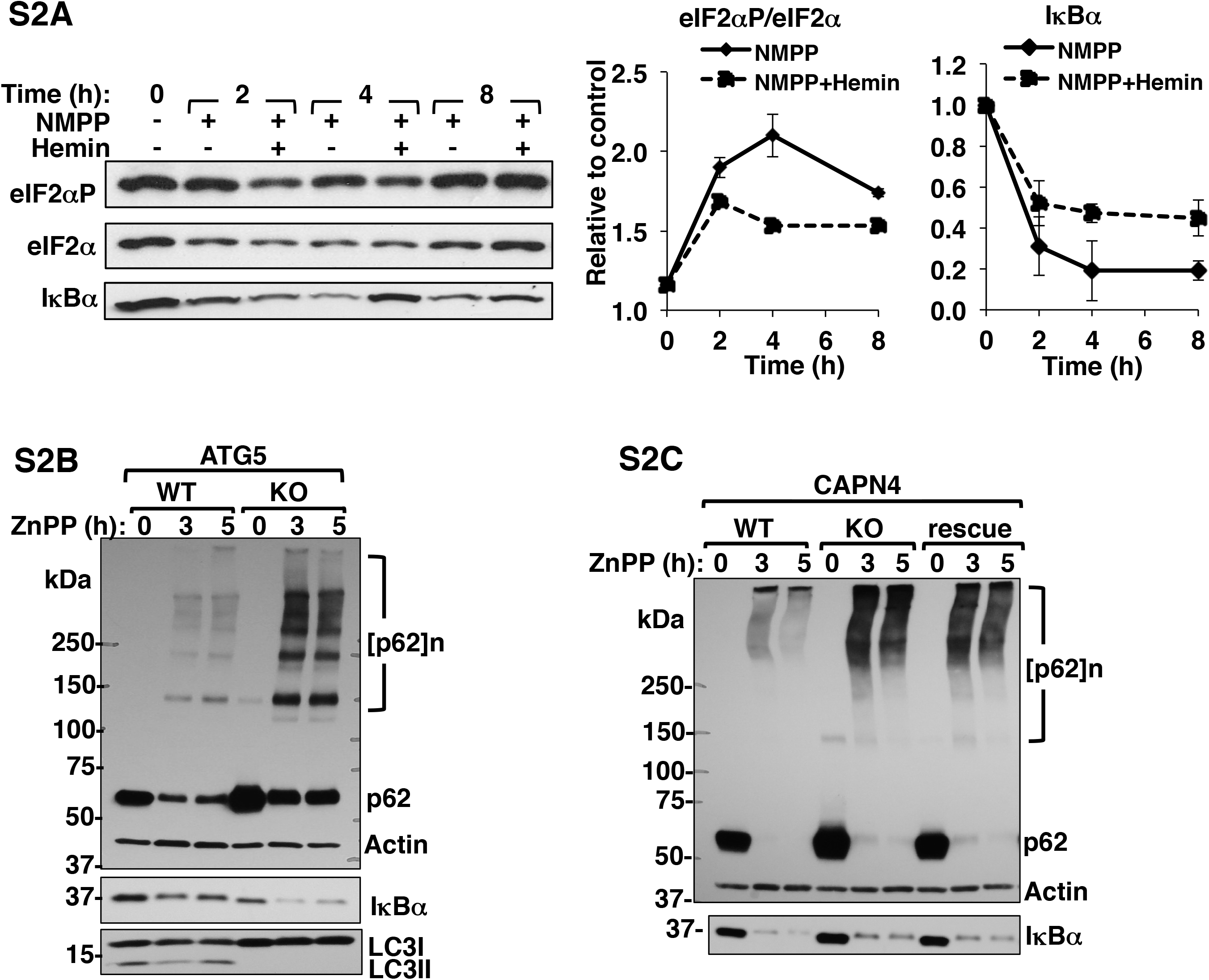
ZnPP-elicited IκBα loss is independent of heme deficiency elicited HRI-activation, autophagy and calpain-mediated proteolysis (Related to Fig 1). **A**. IB analyses of eIF2α, eIF2αP and IκBα in lysates from mouse hepatocytes treated with NMPP or NMPP plus hemin for the indicated times. The temporal profiles of the ratio of eIF2αP to total eIF2α as well as IκBα content were determined (Mean ± SD, n=3). **B**. ATG5 wild-type (ATG5WT) and knockout (ATG5KO) MEF cells were treated with 10 μM ZnPP for the indicated time. Cell lysates were used for IB analysis of p62, IκBα, and LC3, with actin as the loading control. **C**. MEF cells wild-type calpain (capn4WT), capn4 knockout (capn4KO) and capn4KO-rescue (wherein capn4 knockout cells were rescued by transfection of a Capn4 lentiviral vector) were treated with ZnPP for the indicated time. Cell lysates were used for IB analysis of p62 and IκBα with actin as the loading control.

**FIGURE S3.**
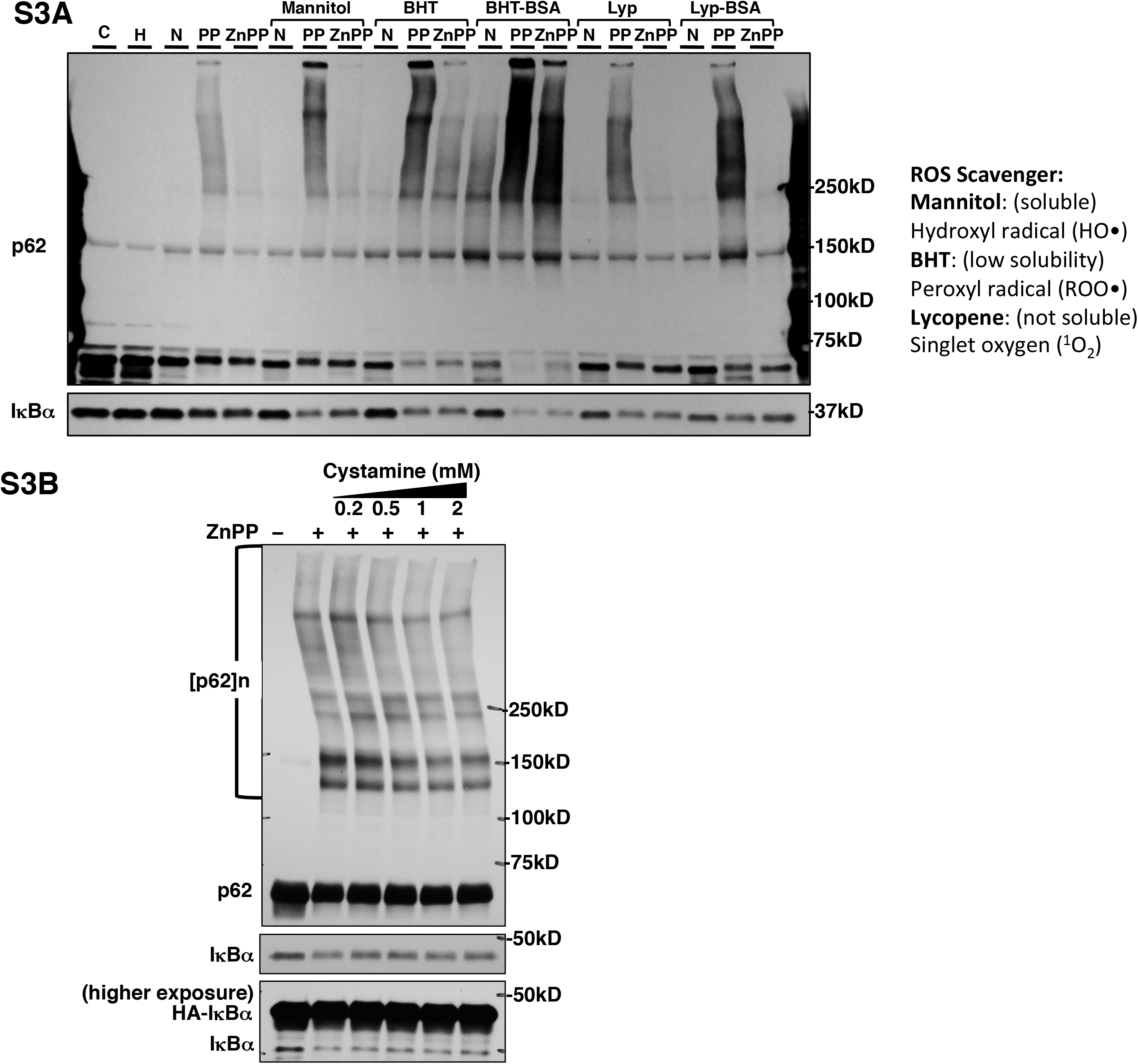
ZnPP-elicited IκBα loss and p62 cannot be reversed by antioxidants or TG2 inhibitors (Related to Fig 1). **A**. Primary mouse hepatocytes were left untreated, or pretreated for 1 h with 50mM mannitol, or 100µM BHT, or 50µM lycopene (Lyp), or BHT coupled with BSA to help keep it solubilized (BHT-BSA), or lycopene coupled with BSA (Lyp-BSA), and then treated with vehicle control (C), 10 μM hemin (H), 30 μM NMPP (N), or 10 μM PPIX (PP), or 10 μM ZnPP (ZnPP) for 4 h as indicated. Cell lysates were used for IB analyses of p62 and IκBα. **B**. HEK293T cells were co-transfected with pcDNA6-p62-myc or pCMV4-3HA-IκBα. 48 h after transfection, cells were pretreated with various concentrations of cystamine as indicated and then treated with 10 μM ZnPP for 2 h. Cell lysates were used for IB analyses of p62 and IκBα.

**FIGURE S4.**
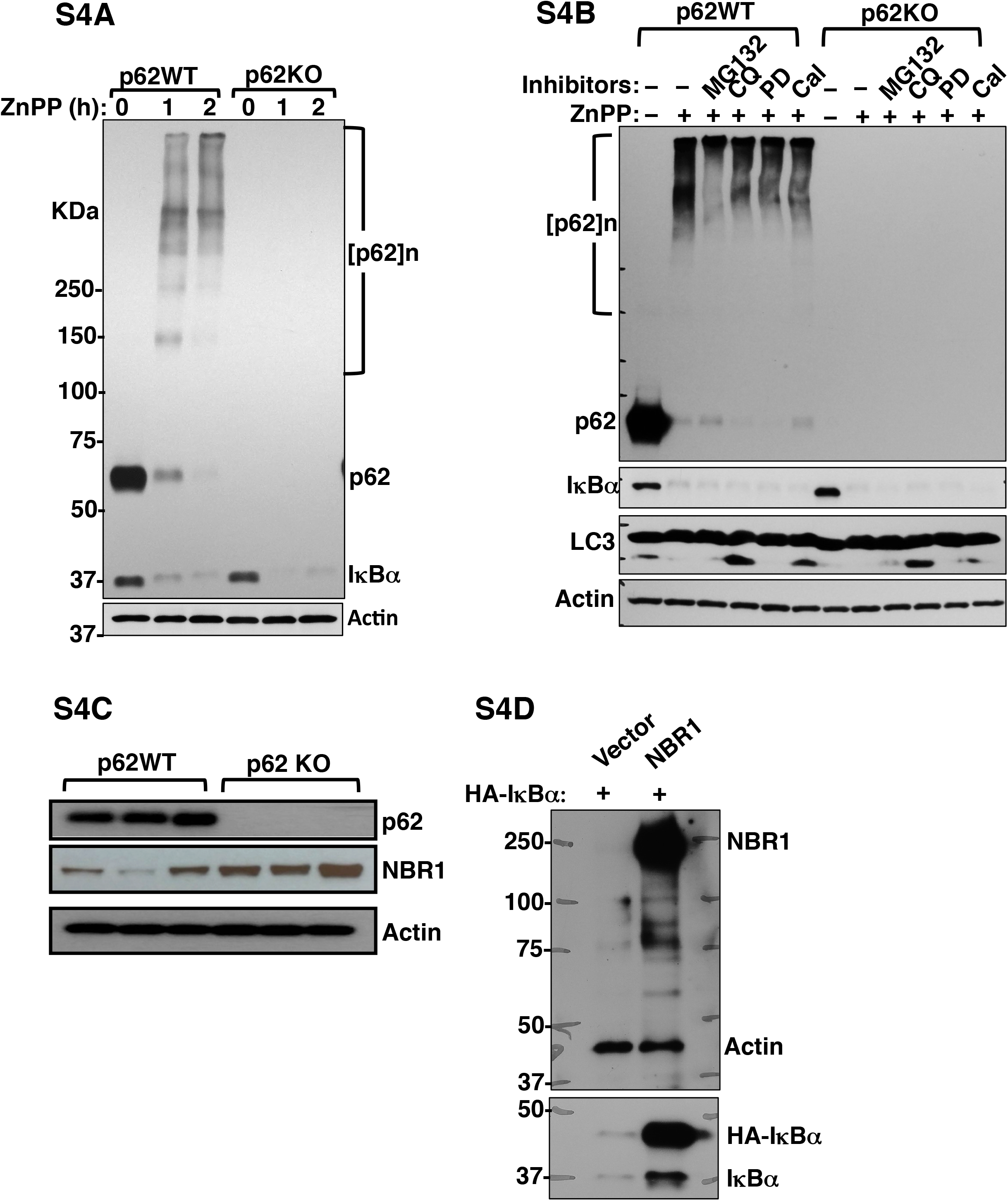
ZnPP-elicited IκBα-loss is independent of NBR1 and other autophagic adapters. (Related to Fig 2). **A.** p62WT and p62KO MEF cells were treated with 10 μM ZnPP for the indicated times. Cell lysates were used for IB analyses of p62 and IkBa, with actin as the loading control. **B.** p62WT and p62KO MEF cells were pretreated for 1 h with inhibitors of various protein degradation pathways: MG132 (20 μM; proteasomal), chloroquine (CQ, 100 μM; lysosomal), PD150606 (PD, 200 μM; calpain1/2), or calpeptin (Cal, 200 μM; cathepsin and calpain), and then treated with 10 μM ZnPP for 2 h. Cell lysates were used for IB analyses. **C**. Lysates from p62WT and p62KO primary mouse hepatocytes were used for IB analyses of p62, and NBR1 with actin as the loading control. The three lanes correspond to p62 WT and p62 KO hepatocytes from three individual mice. **D**. HEK293T cells were co-transfected with pCMV4-3HA-IκBα with either pcDNA3 empty vector or pcDNA3-NBR1 for 48 h. Cell lysates were used for IB analyses of NBR1 and IκBα with actin as the loading control.

**FIGURE S5.**
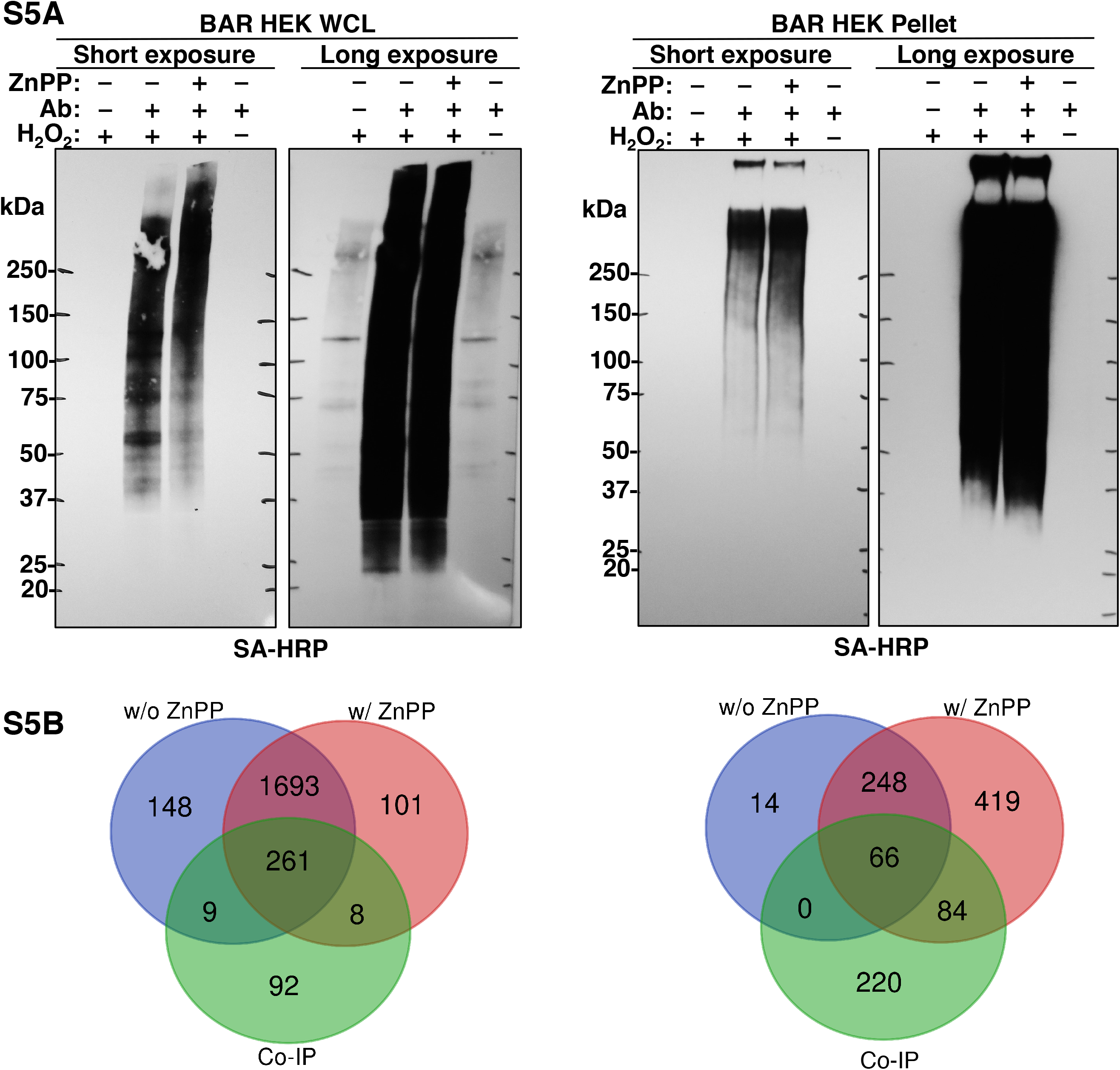
Characterization of biotinylation via IκBα antibody recognition (BAR). (Related to Fig 5B). (**A**) HEK293T cells were transfected with with pCMV4-3HA-IκBα for 48 h. Some cells were treated with 10 μM ZnPP for 2 h. Cells were fixed and then stained with IκBα antibody for downstream HRP antibody recognition dependent biotinylation labeling as described in Methods. Cells were lysed, sub-fractionated into whole cell lysates (WCL) and pellet as described in Methods, separated by SDS-PAGE and analyzed by blotting with streptavidin-HRP. Negative controls in which IκBα antibody in substrate recognition or H_2_O_2_ in labeling reaction was omitted are shown in lanes 1 and 4 of each blot. The band pattern shows that biotinylation is dependent on the presence of both IκBα antibody and H_2_O_2_, suggesting the specificity of biotinylation of endogenous IκBα-proximal proteins. The up-shift of bands in WCL fraction and an increased intensity of bands in the pellet fraction in ZnPP treated cells suggest that IκBα interacts with ZnPP-aggregated proteins. After higher exposure, in the negative control lanes from WCL fraction, some faint bands representing endogenous biotinylated proteins are detected (i.e.130, 75 and 72 kDa). (**B**). Biotinylated proteins within each lysate (including negative controls) were then enriched using streptavidin-coated magnetic beads and trypsin/ Lys-C digested for LC-MS/MS proteomic analyses. Non-specific protein binders were filtered out by comparison to negative controls, and only proteins with an enrichment ratio of ≥ 4 considered specifically biotinylated by BAR. The numbers of proteins identified using BAR in cells treated with or without ZnPP in comparison with the co-IP method in both fractions are shown in the Venn diagrams.

## References

[1] Strnad P, Zatloukal K, Stumptner C, Kulaksiz H, Denk H. Mallory-Denk-bodies: lessons from keratin-containing hepatic inclusion bodies. Biochim Biophys Acta 2008;1782:764–774.

[2] Zatloukal K, Stumptner C, Fuchsbichler A, Heid H, Schnoelzer M, Kenner L, et al. p62 Is a common component of cytoplasmic inclusions in protein aggregation diseases. Am J Pathol 2002;160:255–263.

[3] Jensen K, Gluud C. The Mallory body: morphological, clinical and experimental studies (Part 1 of a literature survey). Hepatology 1994;20:1061–1077.

[4] Denk H, Gschnait F, Wolff K. Hepatocellar hyalin (Mallory bodies) in long term griseofulvin-treated mice: a new experimental model for the study of hyalin formation. Lab Invest 1975;32:773–776.

[5] Yokoo H, Harwood TR, Racker D, Arak S. Experimental production of Mallory bodies in mice by diet containing 3,5-diethoxycarbonyl-1,4-dihydrocollidine. Gastroenterology 1982;83:109–113.

[6] Holley AE, Frater Y, Gibbs AH, De Matteis F, Lamb JH, Farmer PB, et al. Isolation of two N-monosubstituted protoporphyrins, bearing either the whole drug or a methyl group on the pyrrole nitrogen atom, from liver of mice given griseofulvin. Biochem J 1991;274 (Pt 3):843–848.

[7] Ortiz de Montellano PR, Beilan HS, Kunze KL. N-Alkylprotoporphyrin IX formation in 3,5-dicarbethoxy-1,4-dihydrocollidine-treated rats. Transfer of the alkyl group from the substrate to the porphyrin. J Biol Chem 1981;256:6708–6713.

[8] Singla A, Moons DS, Snider NT, Wagenmaker ER, Jayasundera VB, Omary MB. Oxidative stress, Nrf2 and keratin up-regulation associate with Mallory-Denk body formation in mouse erythropoietic protoporphyria. Hepatology 2012;56:322–331.

[9] Schiff ER, Maddrey WC, Sorrell MF. Hepatic Histopathology. Schiff’s Diseases of the Liver: Wiley; 2011. p. 280–282.

[10] Davies R, Schuurman A, Barker CR, Clothier B, Chernova T, Higginson FM, et al. Hepatic gene expression in protoporphyic Fech mice is associated with cholestatic injury but not a marked depletion of the heme regulatory pool. Am J Pathol 2005;166:1041–1053.

[11] Gant TW, Baus PR, Clothier B, Riley J, Davies R, Judah DJ, et al. Gene expression profiles associated with inflammation, fibrosis, and cholestasis in mouse liver after griseofulvin. EHP Toxicogenomics 2003;111:37–43.

[12] Stumptner C, Fuchsbichler A, Heid H, Zatloukal K, Denk H. Mallory body--a disease-associated type of sequestosome. Hepatology 2002;35:1053–1062.

[13] Singla A, Griggs NW, Kwan R, Snider NT, Maitra D, Ernst SA, et al. Lamin aggregation is an early sensor of porphyria-induced liver injury. J Cell Sci 2013;126:3105–3112.

[14] Currais A, Fischer W, Maher P, Schubert D. Intraneuronal protein aggregation as a trigger for inflammation and neurodegeneration in the aging brain. FASEB J 2017;31:5–10.

[15] Nivon M, Fort L, Muller P, Richet E, Simon S, Guey B, et al. NFkappaB is a central regulator of protein quality control in response to protein aggregation stresses via autophagy modulation. Mol Biol Cell 2016;27:1712–1727.

[16] Liou HC, Baltimore D. Regulation of the NF-kappa B/rel transcription factor and I kappa B inhibitor system. Curr Opin Cell Biol 1993;5:477–487.

[17] He G, Karin M. NF-kappaB and STAT3 - key players in liver inflammation and cancer. Cell Res 2011;21:159–168.

[18] Azimifar SB, Nagaraj N, Cox J, Mann M. Cell-type-resolved quantitative proteomics of murine liver. Cell Metab 2014;20:1076–1087.

[19] Han Y, Brasier AR. Mechanism for biphasic rel A. NF-kappaB1 nuclear translocation in tumor necrosis factor alpha-stimulated hepatocytes. J Biol Chem 1997;272:9825–9832.

[20] Rao P, Hayden MS, Long M, Scott ML, West AP, Zhang D, et al. IkappaBbeta acts to inhibit and activate gene expression during the inflammatory response. Nature 2010;466:1115–1119.

[21] Wisniewski JR, Vildhede A, Noren A, Artursson P. In-depth quantitative analysis and comparison of the human hepatocyte and hepatoma cell line HepG2 proteomes. J Proteomics 2016;136:234–247.

[22] Beg AA, Finco TS, Nantermet PV, Baldwin AS, Jr. Tumor necrosis factor and interleukin-1 lead to phosphorylation and loss of I kappa B alpha: a mechanism for NF-kappa B activation. Mol Cell Biol 1993;13:3301–3310.

[23] Traenckner EB, Wilk S, Baeuerle PA. A proteasome inhibitor prevents activation of NF-kappa B and stabilizes a newly phosphorylated form of I kappa B-alpha that is still bound to NF-kappa B. EMBO J 1994;13:5433–5441.

[24] Weil R, Laurent-Winter C, Israel A. Regulation of IkappaBbeta degradation. Similarities to and differences from IkappaBalpha. J Biol Chem 1997;272:9942–9949.

[25] Arenzana-Seisdedos F, Thompson J, Rodriguez MS, Bachelerie F, Thomas D, Hay RT. Inducible nuclear expression of newly synthesized I kappa B alpha negatively regulates DNA-binding and transcriptional activities of NF-kappa B. Mol Cell Biol 1995;15:2689–2696.

[26] Thompson JE, Phillips RJ, Erdjument-Bromage H, Tempst P, Ghosh S. I kappa B-beta regulates the persistent response in a biphasic activation of NF-kappa B. Cell 1995;80:573–582.

[27] Arenzana-Seisdedos F, Turpin P, Rodriguez M, Thomas D, Hay RT, Virelizier JL, et al. Nuclear localization of I kappa B alpha promotes active transport of NF-kappa B from the nucleus to the cytoplasm. J Cell Sci 1997;110 (Pt 3):369–378.

[28] Cox TM. Protoporphyria. In: Karl M. Kadish KMS, Roger Guilard, editor. The Porphyrin Handbook: Medical aspects of porphyrins. USA: Elsevier science; 2003.

[29] Thapar M, Bonkovsky HL. The diagnosis and management of erythropoietic protoporphyria. Gastroenterol Hepatol (N Y) 2008;4:561–566.

[30] Balwani M, Doheny D, Bishop DF, Nazarenko I, Yasuda M, Dailey HA, et al. Loss-of-function ferrochelatase and gain-of-function erythroid-specific 5-aminolevulinate synthase mutations causing erythropoietic protoporphyria and x-linked protoporphyria in North American patients reveal novel mutations and a high prevalence of X-linked protoporphyria. Mol Med 2013;19:26–35.

[31] Chen JJ. Regulation of protein synthesis by the heme-regulated eIF2alpha kinase: relevance to anemias. Blood 2007;109:2693–2699.

[32] Cuervo AM, Hu W, Lim B, Dice JF. IkappaB is a substrate for a selective pathway of lysosomal proteolysis. Mol Biol Cell 1998;9:1995–2010.

[33] Chen F, Lu Y, Kuhn DC, Maki M, Shi X, Sun SC, et al. Calpain contributes to silica-induced I kappa B-alpha degradation and nuclear factor-kappa B activation. Arch Biochem Biophys 1997;342:383–388.

[34] Kirkin V, Lamark T, Sou YS, Bjorkoy G, Nunn JL, Bruun JA, et al. A role for NBR1 in autophagosomal degradation of ubiquitinated substrates. Mol Cell 2009;33:505–516.

[35] Lazarou M, Sliter DA, Kane LA, Sarraf SA, Wang C, Burman JL, et al. The ubiquitin kinase PINK1 recruits autophagy receptors to induce mitophagy. Nature 2015;524:309–314.

[36] Maitra D, Elenbaas JS, Whitesall SE, Basrur V, D’Alecy LG, Omary MB. Ambient Light Promotes Selective Subcellular Proteotoxicity after Endogenous and Exogenous Porphyrinogenic Stress. J Biol Chem 2015;290:23711–23724.

[37] Elenbaas JS, Maitra D, Liu Y, Lentz SI, Nelson B, Hoenerhoff MJ, et al. A precursor-inducible zebrafish model of acute protoporphyria with hepatic protein aggregation and multiorganelle stress. FASEB J 2016;30:1798–1810.

[38] Bar DZ, Atkatsh K, Tavarez U, Erdos MR, Gruenbaum Y, Collins FS. Biotinylation by antibody recognition-a method for proximity labeling. Nat Methods 2018;15:127–133.

[39] Klement JF, Rice NR, Car BD, Abbondanzo SJ, Powers GD, Bhatt PH, et al. IkappaBalpha deficiency results in a sustained NF-kappaB response and severe widespread dermatitis in mice. Mol Cell Biol 1996;16:2341–2349.

[40] Tergaonkar V, Correa RG, Ikawa M, Verma IM. Distinct roles of IkappaB proteins in regulating constitutive NF-kappaB activity. Nat Cell Biol 2005;7:921–923.

[41] Yang H, Hu HY. Sequestration of cellular interacting partners by protein aggregates: implication in a loss-of-function pathology. FEBS J 2016;283:3705–3717.

[42] Lahiri P, Schmidt V, Smole C, Kufferath I, Denk H, Strnad P, et al. p62/Sequestosome-1 Is Indispensable for Maturation and Stabilization of Mallory-Denk Bodies. PLoS One 2016;11:e0161083.

[43] Olzscha H, Schermann SM, Woerner AC, Pinkert S, Hecht MH, Tartaglia GG, et al. Amyloid-like aggregates sequester numerous metastable proteins with essential cellular functions. Cell 2011;144:67–78.

[44] Ferreiro DU, Komives EA. Molecular mechanisms of system control of NF-kappaB signaling by IkappaBalpha. Biochemistry 2010;49:1560–1567.

[45] Ghosh S, Karin M. Missing pieces in the NF-kappaB puzzle. Cell 2002;109 Suppl:S81–96.

[46] Fagerlund R, Kinnunen L, Kohler M, Julkunen I, Melen K. NF-{kappa}B is transported into the nucleus by importin {alpha}3 and importin {alpha}4. J Biol Chem 2005;280:15942–15951.

[47] Lu M, Zak J, Chen S, Sanchez-Pulido L, Severson DT, Endicott J, et al. A code for RanGDP binding in ankyrin repeats defines a nuclear import pathway. Cell 2014;157:1130–1145.

[48] Ball JR, Ullman KS. Versatility at the nuclear pore complex: lessons learned from the nucleoporin Nup153. Chromosoma 2005;114:319–330.

[49] Nakielny S, Shaikh S, Burke B, Dreyfuss G. Nup153 is an M9-containing mobile nucleoporin with a novel Ran-binding domain. EMBO J 1999;18:1982–1995.

[50] Partridge JR, Schwartz TU. Crystallographic and biochemical analysis of the Ran-binding zinc finger domain. J Mol Biol 2009;391:375–389.

[51] Walde S, Thakar K, Hutten S, Spillner C, Nath A, Rothbauer U, et al. The nucleoporin Nup358/RanBP2 promotes nuclear import in a cargo- and transport receptor-specific manner. Traffic 2012;13:218–233.

[52] Desterro JM, Rodriguez MS, Hay RT. SUMO-1 modification of IkappaBalpha inhibits NF-kappaB activation. Mol Cell 1998;2:233–239.

[53] Ofengeim D, Ito Y, Najafov A, Zhang Y, Shan B, DeWitt JP, et al. Activation of necroptosis in multiple sclerosis. Cell Rep 2015;10:1836–1849.

[54] Boeynaems S, Bogaert E, Van Damme P, Van Den Bosch L. Inside out: the role of nucleocytoplasmic transport in ALS and FTLD. Acta Neuropathol 2016;132:159–173.

[55] Nakahara J, Kanekura K, Nawa M, Aiso S, Suzuki N. Abnormal expression of TIP30 and arrested nucleocytoplasmic transport within oligodendrocyte precursor cells in multiple sclerosis. J Clin Invest 2009;119:169–181.

[56] Yamashita T, Aizawa H, Teramoto S, Akamatsu M, Kwak S. Calpain-dependent disruption of nucleo-cytoplasmic transport in ALS motor neurons. Sci Rep 2017;7:39994.

[57] Tutois S, Montagutelli X, Da Silva V, Jouault H, Rouyer-Fessard P, Leroy-Viard K, et al. Erythropoietic protoporphyria in the house mouse. A recessive inherited ferrochelatase deficiency with anemia, photosensitivity, and liver disease. J Clin Invest 1991;88:1730–1736.

[58] Dubbelman TM, de Goeij AF, van Steveninck J. Photodynamic effects of protoporphyrin on human erythrocytes. Nature of the cross-linking of membrane proteins. Biochim Biophys Acta 1978;511:141–151.

[59] Afonso SG, Enriquez de Salamanca R, Batlle AM. The photodynamic and non-photodynamic actions of porphyrins. Braz J Med Biol Res 1999;32:255–266.

## References

[1] Jiang HY, Wek SA, McGrath BC, Scheuner D, Kaufman RJ, Cavener DR, et al. Phosphorylation of the alpha subunit of eukaryotic initiation factor 2 is required for activation of NF-kappaB in response to diverse cellular stresses. Mol Cell Biol 2003;23:5651–5663.

[2] Deng J, Lu PD, Zhang Y, Scheuner D, Kaufman RJ, Sonenberg N, et al. Translational repression mediates activation of nuclear factor kappa B by phosphorylated translation initiation factor 2. Mol Cell Biol 2004;24:10161–10168.

[3] Chen JJ. Regulation of protein synthesis by the heme-regulated eIF2alpha kinase: relevance to anemias. Blood 2007;109:2693–2699.

[4] Cuervo AM, Hu W, Lim B, Dice JF. IkappaB is a substrate for a selective pathway of lysosomal proteolysis. Mol Biol Cell 1998;9:1995–2010.

[5] Pankiv S, Clausen TH, Lamark T, Brech A, Bruun JA, Outzen H, et al. p62/SQSTM1 binds directly to Atg8/LC3 to facilitate degradation of ubiquitinated protein aggregates by autophagy. J Biol Chem 2007;282:24131–24145.

[6] Chen F, Lu Y, Kuhn DC, Maki M, Shi X, Sun SC, et al. Calpain contributes to silica-induced I kappa B-alpha degradation and nuclear factor-kappa B activation. Arch Biochem Biophys 1997;342:383–388.

[7] Han Y, Weinman S, Boldogh I, Walker RK, Brasier AR. Tumor necrosis factor-alpha-inducible IkappaBalpha proteolysis mediated by cytosolic m-calpain. A mechanism parallel to the ubiquitin-proteasome pathway for nuclear factor-kappab activation. J Biol Chem 1999;274:787–794.

[8] Cox TM. Protoporphyria. In: Karl M. Kadish KMS, Roger Guilard, editor. The Porphyrin Handbook: Medical aspects of porphyrins. USA: Elsevier science; 2003.

[9] Drummond GS, Kappas A. Prevention of neonatal hyperbilirubinemia by tin protoporphyrin IX, a potent competitive inhibitor of heme oxidation. Proc Natl Acad Sci U S A 1981;78:6466–6470.

[10] Lamola AA, Yamane T. Zinc protoporphyrin in the erythrocytes of patients with lead intoxication and iron deficiency anemia. Science 1974;186:936–938.

[11] Freesemann AG, Gross U, Bensidhoum M, de Verneuil H, Doss MO. Immunological, enzymatic and biochemical studies of uroporphyrinogen III-synthase deficiency in 20 patients with congenital erythropoietic porphyria. Eur J Biochem 1998;257:149–153.

[12] To-Figueras J, Millet O, Herrero C. Congenital Erythropoietic Porphyria. Handbook of Porphyrin Science (Volume 29); 2013. p. 151–217.

[13] Han XM, Lee G, Hefner C, Maher JJ, Correia MA. Heme-reversible impairment of CYP2B1/2 induction in heme-depleted rat hepatocytes in primary culture: translational control by a hepatic alpha-subunit of the eukaryotic initiation factor kinase? J Pharmacol Exp Ther 2005;314:128–138.

[14] Chisolm J, Jr., Brown DH. Micro-scale photofluorometric determination of “free erythrocyte pophyrin” (protoporphyrin IX). Clin Chem 1975;21:1669–1682.

[15] Ku NO, Toivola DM, Zhou Q, Tao GZ, Zhong B, Omary MB. Studying simple epithelial keratins in cells and tissues. Methods Cell Biol 2004;78:489–517.

[16] Che Y, Khavari PA. Research Techniques Made Simple: Emerging Methods to Elucidate Protein Interactions through Spatial Proximity. The Journal of investigative dermatology 2017;137:e197–e203.

[17] Bar DZ, Atkatsh K, Tavarez U, Erdos MR, Gruenbaum Y, Collins FS. Biotinylation by antibody recognition-a method for proximity labeling. Nat Methods 2018;15:127–133.

[18] Hung V, Udeshi ND, Lam SS, Loh KH, Cox KJ, Pedram K, et al. Spatially resolved proteomic mapping in living cells with the engineered peroxidase APEX2. Nat Protoc 2016;11:456–475.

[19] Rosenfeld J, Capdevielle J, Guillemot JC, Ferrara P. In-gel digestion of proteins for internal sequence analysis after one- or two-dimensional gel electrophoresis. Anal Biochem 1992;203:173–179.

[20] Hellman U, Wernstedt C, Gonez J, Heldin CH. Improvement of an “In-Gel” digestion procedure for the micropreparation of internal protein fragments for amino acid sequencing. Anal Biochem 1995;224:451–455.

[21] Chalkley RJ, Baker PR, Medzihradszky KF, Lynn AJ, Burlingame AL. In-depth analysis of tandem mass spectrometry data from disparate instrument types. Mol Cell Proteomics 2008;7:2386–2398.

[22] Elias JE, Gygi SP. Target-decoy search strategy for increased confidence in large-scale protein identifications by mass spectrometry. Nat Methods 2007;4:207–214.

[23] Ma H, McLean JR, Chao LF, Mana-Capelli S, Paramasivam M, Hagstrom KA, et al. A highly efficient multifunctional tandem affinity purification approach applicable to diverse organisms. Mol Cell Proteomics 2012;11:501–511.

[24] Szklarczyk D, Franceschini A, Wyder S, Forslund K, Heller D, Huerta-Cepas J, et al. STRING v10: protein-protein interaction networks, integrated over the tree of life. Nucleic Acids Res 2015;43:D447–452.

[25] Shannon P, Markiel A, Ozier O, Baliga NS, Wang JT, Ramage D, et al. Cytoscape: a software environment for integrated models of biomolecular interaction networks. Genome Res 2003;13:2498–2504.

[26] Kyte J, Doolittle RF. A simple method for displaying the hydropathic character of a protein. J Mol Biol 1982;157:105–132.

[27] Peng Z, Mizianty MJ, Kurgan L. Genome-scale prediction of proteins with long intrinsically disordered regions. Proteins 2014;82:145–158.

